# Transposable Element Derepression-mediated IFN-I Signaling Enhances Stem/Progenitor Cell Activity After Irradiation

**DOI:** 10.1101/2024.02.14.580306

**Authors:** Davide Cinat, Ryan van der Wal, Mirjam Baanstra, Abel Soto-Gamez, Anne L. Jellema-de Bruin, Uilke Brouwer, Marc-Jan van Goethem, Marcel A.T.M. van Vugt, Lara Barazzuol, Rob P. Coppes

## Abstract

Radiotherapy is a mainstay in cancer treatment, aiming to maximize DNA damage in tumors while minimizing harm to surrounding healthy tissues. However, the collateral damage to normal tissues, especially stem/progenitor cells essential for tissue regeneration and organ function, remains a significant challenge. Here, we investigate the molecular responses to photon and proton irradiation, two key modalities in head and neck cancer treatment, using organoids. Multiomics analysis reveals a stronger double-stranded RNA (dsRNA)-induced type I interferon (IFN-I) response following proton irradiation, driven by loss of heterochromatin regulators and derepression of transposable elements (TEs). This response, mediated by the cytoplasmic sensor RIG-I, enhances the inflammatory signaling initiated by the canonical dsDNA sensors cGAS and ZBP1. Genetic and pharmacological modulation of IFN-I signaling *in vitro* and *in vivo* demonstrates its critical role in enhancing stem/progenitor cell activity post-irradiation. Our findings reveal a pro-regenerative role of TE derepression-mediated IFN-I response suggesting this pathway as a promising therapeutic target to mitigate radiation-induced side effects.

**Teaser:** Transposable element-mediated type I interferon signaling enhances stem/progenitor cell activity after irradiation.

## Introduction

The DNA damage-induced response is an intricate process culminating in the induction of inflammatory and immune reactions (*1*, *2*). However, DNA damage-induced inflammation has predominantly been studied in tumor models (*3*, *4*), overlooking its potential significance in non-transformed tissues that also undergo DNA damage during cancer treatment. This is particularly relevant in the context of head and neck cancer patients, where healthy salivary glands are often unavoidably exposed to radiation (*5*). Such exposure can lead to normal tissue toxicity, loss of salivary gland function, and ultimately, radiation-induced xerostomia with devastating consequences for the patients’ quality of life (*6*).

Adult normal tissue stem cells are crucial for regenerating damaged tissues (*7*). Our previous work demonstrated that sparing the salivary gland region containing stem/progenitor cells significantly reduced the risk of post-radiotherapy xerostomia (*8*, *9*). However, stem cell proliferation and self-renewal capacity can also be influenced by stress-induced microenvironmental changes (*1*, *10*, *11*). For instance, the expression of type I interferons (IFN-I), can be triggered by radiation-induced DNA damage (*12*, *13*), leading to the activation of an immune response. However, the exact mechanism underlying the induction of IFN-I expression and its subsequent impact on normal tissue stem cells remains largely unknown.

Due to their scarcity and often quiescent state (*14*), studying the mechanisms regulating salivary gland stem cells *in vivo* is challenging. For this reason, stem cell-derived salivary gland organoids have been instrumental in elucidating the signaling pathways that regulate stem/progenitor self-renewal and differentiation *in vitro* (*15–18*). Moreover, these organoids closely resemble the normal tissue of origin (*19*) and have advanced our understanding of the mechanisms triggered by radiation-induced DNA damage (*18*, *20*).

Importantly, the normal tissue response may vary depending on the complexity of DNA damage induced by different radiation modalities, as evidenced in the context of proton therapy compared to conventional photon-based radiotherapy (*5*). Protons are characterized by physical properties that allow for a more precise radiation dose deposition to the tumor while minimizing exposure to surrounding normal tissues (*5*). Furthermore, recent studies have revealed that proton irradiation can elicit distinct DNA damage responses and activate specific pathways (*21–23*), leading to more effective killing of cancer cells (*21*, *24*).

However, the understanding of the molecular effects of irradiation on normal tissue and its stem cells remains largely unexplored. In this regard, we used mouse salivary gland organoids enriched in stem and progenitor cells (*16*) as a model to investigate the response of normal tissue to photon and proton irradiation. We show that proton irradiation elicits a higher IFN-I response via derepression of TEs and cytoplasmic dsRNA formation, ultimately enhancing stem/progenitor cell activity and self-renewal capacity both in organoids and mouse models.

## Results

### Photon and proton irradiation lead to pronounced IFN-I signaling activation

To better understand the effects of proton and photon irradiation, two clinically relevant treatment modalities for head and neck cancer, we irradiated organoids on day 5 with 7 Gy photons or protons and assessed their survival and self-renewal capacity 6 days after irradiation (day 11) (Fig. 1A). Based on a previous study, this dose and timepoint replicated the cytotoxicity, regenerative responses, and mechanistic changes observed *in vivo* (*20*). Surprisingly, despite the similar decrease in organoid forming efficiency (OFE) observed after photon and proton irradiation at day 11 (fig. S1, A and B), organoids irradiated with 7 Gy protons exhibited a higher self-renewal capacity compared to those irradiated with 7 Gy photons. This was measured as OFE at day 18 (one week after passaging) per 10,000 seeded cells on day 11 (Fig. 1, B and C), suggesting a differential response in the stem/progenitor cell compartment of these organoids. To investigate the mechanisms underlying this difference, we performed bulk RNA-sequencing (RNA-seq) on organoids 6 days after irradiation (fig. S1C). Bioinformatic analysis revealed profound radiation-induced transcriptional changes, involving a strong, dose-dependent upregulation of genes associated with immune response and inflammation (Fig. 1D and fig. S1D). Particularly, the interferon-beta (IFN-β) response appeared as one of the most enriched biological processes after irradiation (Fig. 1D and fig. S1E) with *Ifnb* and several IFN-stimulated genes (ISGs) showing higher expression after proton compared to photon irradiation (Fig. 1, E and F and fig. S1, F to H). In line with these findings, higher protein levels of STAT1 and p-STAT1 were measured after proton irradiation, indicating a more pronounced activation of IFN-I/STAT1/ISGs signaling compared to photon (Fig. 1, G and H and fig. S1I).

**Figure 1:**
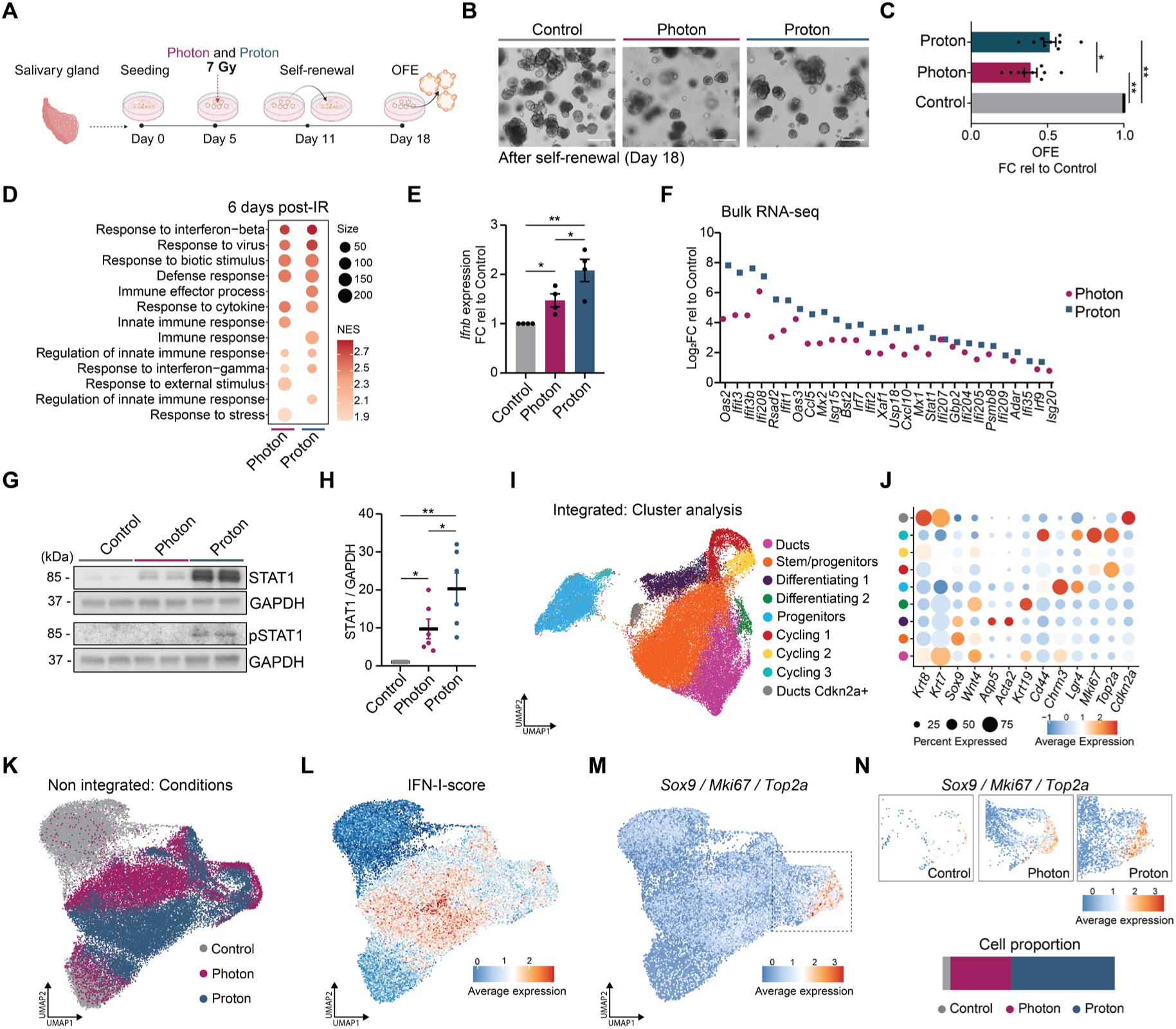
Proton irradiation leads to a higher IFN-I-signaling upregulation and stem/progenitor cell activity. (**A**) Schematic representation of the self-renewal assay. Organoids were irradiated at day 5 and harvested at day 11 (6 days post irradiation). Organoids were dissociated into single cells and re-plated to assess their self-renewal capacity at day 18, shown as percentage of organoid forming efficiency (OFE%). (**B**) Representative images of organoids in culture after self-renewal (day 18). Scale bar, 100 µm. (**C**) Organoid quantification shown as OFE fold change (FC) relative to control samples (means ± s.e.m; n = 9 animals/condition). One-way ANOVA with Tukey’s multiple comparison test. (**D**) Top 10 significant (p.adj. < 0.05) biological processes in proton vs control and photon vs control at 6 days post irradiation. (**E**) rt-qPCR analysis of *Ifnb* at 6 days post irradiation. Data is shown as FC relative to control samples (means ± s.e.m; n = 4 animals/condition). One-way ANOVA with Tukey’s multiple comparison test. (**F**) Gene expression level of significant (p < 0.05) ISGs extrapolated from the bulk RNA-seq data of organoids at 6 days post irradiation. Data is shown as Log_2_FC relative to control samples. (**G**) Western blot analysis of STAT1, pSTAT1 and GAPDH of organoids at 6 days post irradiation. All biological replicates are shown in fig. S1I. (**H**) Western blot quantification of STAT1 (Fig. 1G and fig. S1I). STAT1 protein levels were normalized for GAPDH. Data is shown as FC relative to control samples (means ± s.e.m; n = 6 animals/condition). One-way ANOVA with Tukey’s multiple comparison test. (**I**) UMAP of the integrated dataset of organoids at 6 days post irradiation (day 11) showing the main cell populations. (**J**) Dot plot showing representative cell type marker genes for each population, cluster colors are the same as in Figure 1I. (**K**) UMAP of the non-integrated dataset of organoids at 6 days post irradiation (day 11) showing the different conditions. (**L**) UMAP of the non-integrated dataset of organoids at 6 days post irradiation (day 11) showing the average expression of the IFN-I-related genes *Stat1, Ifi27l2a, Ifih1, Ifi44, Ifit1* and *Irf9* (shown as IFN-I-score). (**M**) UMAP of the non-integrated dataset of organoids at 6 days post irradiation (day 11) showing the average expression of *Sox9, Mki67* and *Top2a.* (**N**) Zoom-ins of the UMAP in Fig. 1M showing the average expression of *Sox9, Mki67* and *Top2a* (top) and their proportion in control, photon and proton (bottom).

Recent studies have highlighted IFN-I signaling as a key regulator of stem cell activity and tissue regeneration (*25–28*). Given that previous characterization of this salivary gland organoid model identified an enrichment of distinct salivary stem/progenitor cell populations (*18*, *29*), we performed single-cell RNA sequencing (scRNA-seq) on control, photon-irradiated, and proton-irradiated organoids at day 11 to assess potential differences in these rare cell populations at single-cell level. Analysis of the integrated dataset (Fig. 1I and fig. S2, A and B) revealed distinct cell populations, including ductal cells expressing *Krt7*, *Krt8*, and *Krt19*; two stem/progenitor cell populations expressing *Sox9*, *Wnt4* and *Cd44, Lgr4*; differentiating cells expressing *Aqp5, Acta2*, and *Krt19*; cycling cells marked by *Mki67* and *Top2a*; and cell cycle-arrested duct cells expressing *Krt8, Krt7*, and *Cdkn2a* (Fig. 1J).

In line with the bulk RNA-seq data (Fig. 1, D and F), irradiation led to an elevated IFN-I response, which was more pronounced in proton-irradiated cells compared to photon-irradiated cells (Fig. 1, K and L). While all cell populations exhibited a robust IFN-I signature (fig. S2C), *Sox9*-expressing cells displayed the strongest overlap with this response (fig. S2D). Recently, we demonstrated that *Sox9*+ stem/progenitor cells possess stem cell-like properties and the capacity to form organoids (*18*). Further scRNA-seq analysis identified a subset of *Sox9*+ cells enriched with proliferative markers (*Mki67* and *Top2a*), which were more prevalent in irradiated samples compared to control (Fig. 1M, N). Although this population was present in both photon- and proton-irradiated samples (fig. S2B), differential gene expression and gene ontology (GO) term analyses within this cluster revealed a stronger upregulation of processes related to mitotic cell cycle and cell cycle G2/M phase transition upon proton irradiation (fig. S2E). This suggests increased cellular activity, aligning with the higher self-renewal capacity observed after proton irradiation (Fig. 1C). Together, these findings indicate that the IFN-I response triggered by irradiation may positively influence the activity and proliferation of salivary gland stem/progenitor cells, potentially explaining the enhanced self-renewal capacity observed following proton irradiation.

### cGAS and ZBP1 are central regulators of interferon signaling after irradiation

To investigate the molecular mechanisms driving the differential IFN-I signaling observed after photon and proton irradiation, we examined the role of cytoplasmic nucleic acid sensors in mediating this response. The cGAS-STING pathway, which is activated by cytoplasmic dsDNA, is a well-established initiator of IFN-I signaling, typically triggered by DNA damage-induced micronuclei formation at early timepoints (*30*). Activation of this pathway could potentially explain the differential inflammatory response observed after photon and proton irradiation. However, immunofluorescence staining of organoid sections and 2D salivary gland cells revealed a similar increase in cGAS+ micronuclei 2 days after 7 Gy photon and proton irradiation, followed by a decline at later timepoints (Fig. 2, A and B and fig. S2F). Western blot analysis of cytoplasmic fractions (fig. S3A) further confirmed comparable cGAS levels at 2 days post-irradiation (Fig. 2C). Additionally, bulk RNA-seq analysis showed a similar enrichment of immune and inflammatory-related processes (Fig. 2D), along with upregulation of *Ifnb* and ISGs (fig. S3B), suggesting a similar involvement of the cGAS-STING pathway in IFN-I signaling activation across both radiation types. Interestingly, despite the lower levels of cGAS observed 6 days post-irradiation (Fig. 2C), cGAS depletion (fig. S3C) or treatment with the covalent STING inhibitor C-176 significantly reduced ISG expression (Fig. 3, D and F), confirming the critical upstream role of cGAS-STING in regulating the global IFN-I response (*31*).

**Figure 2:**
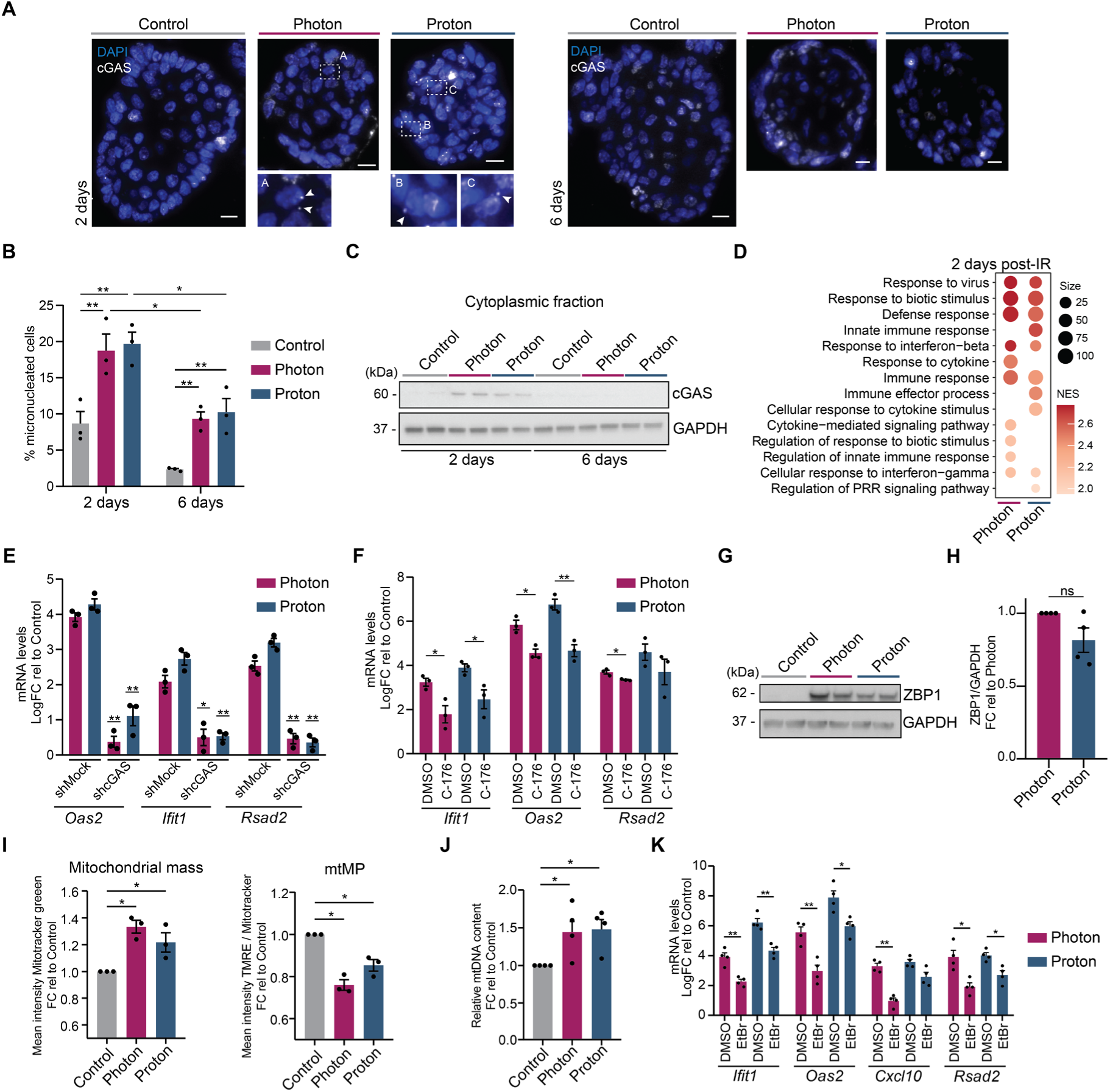
Irradiation-induced IFN-I signaling depends on cGAS and ZBP1. (**A**) Representative images of immunofluorescence staining of organoids sections at 2 days (left) and 6 days (right) post irradiation. Zoom-ins and arrows show cGAS+ micronuclei. Scale bar, 5 µm. (**B**) Quantification of cGAS+ micronucleated cells at 2 and 6 days. Data is shown as percentage (%) of cGAS+ micronucleated cells/total number of cells (means ± s.e.m; n = 3 animals/condition). Two-way ANOVA with Tukey’s multiple comparison test. (**C**) Western blot analysis of cGAS and GAPDH of cytoplasmic fractions extracted from organoids at 2 days and 6 days post irradiation. Validation of the fractionation is shown in fig. S3A. (**D**) Top 10 significant (p.adj. < 0.05) biological processes in proton vs control and photon vs control at 2 days post irradiation. (**E**) rt-qPCR analysis of ISGs in shMock and shcGAS-treated organoids at 6 days post irradiation. Data is shown as Log_2_FC relative to shMock control (means ± s.e.m; n = 3 animals/condition). One-way ANOVA with Tukey’s multiple comparison test. *p and **p are relative to shMock. (**F**) rt-qPCR analysis of ISGs in DMSO and C-176-treated organoids at 6 days post irradiation. Data is shown as Log_2_FC relative to DMSO control (means ± s.e.m; n = 3 animals/condition). Two-sided unpaired *t*-test. (**G**) Western blot analysis of ZBP1 and GAPDH of organoids at 6 days post irradiation. All biological replicates are shown in fig. S3E. (**H**) Western blot quantification of ZBP1 (Fig. 2G and fig. S3E). ZBP1 protein levels were normalized for GAPDH. Data is shown as FC relative to photon samples (means ± s.e.m; n = 4 animals/condition). Two-sided unpaired *t*-test. (**I**) Flow cytometry analysis showing the mitochondrial mass (left panel) and mitochondrial membrane potential (mtMP) (right panel) at 6 days post irradiation. Data is shown as FC relative to control (means ± s.e.m; n = 3 animals/condition). One-way ANOVA with Tukey’s multiple comparison test. (**J**) rt-qPCR analysis of mtDNA genes of cytoplasmic fractions of organoids at 6 days post irradiation. Data is shown as mean expression of *mt16S, mtCOX2, mtDloop3*. Data is shown as FC relative to control samples (means ± s.e.m; n = 4 animals/condition). One-way ANOVA with Tukey’s multiple comparison test. (**K**) rt-qPCR analysis of ISGs in DMSO and EtBr-treated organoids at 6 days post irradiation. Data is shown as Log_2_FC relative to Control (means ± s.e.m; n = 4 animals/condition). Two-sided unpaired *t*-test. In all figures *p < 0.05, **p < 0.01.

**Figure 3:**
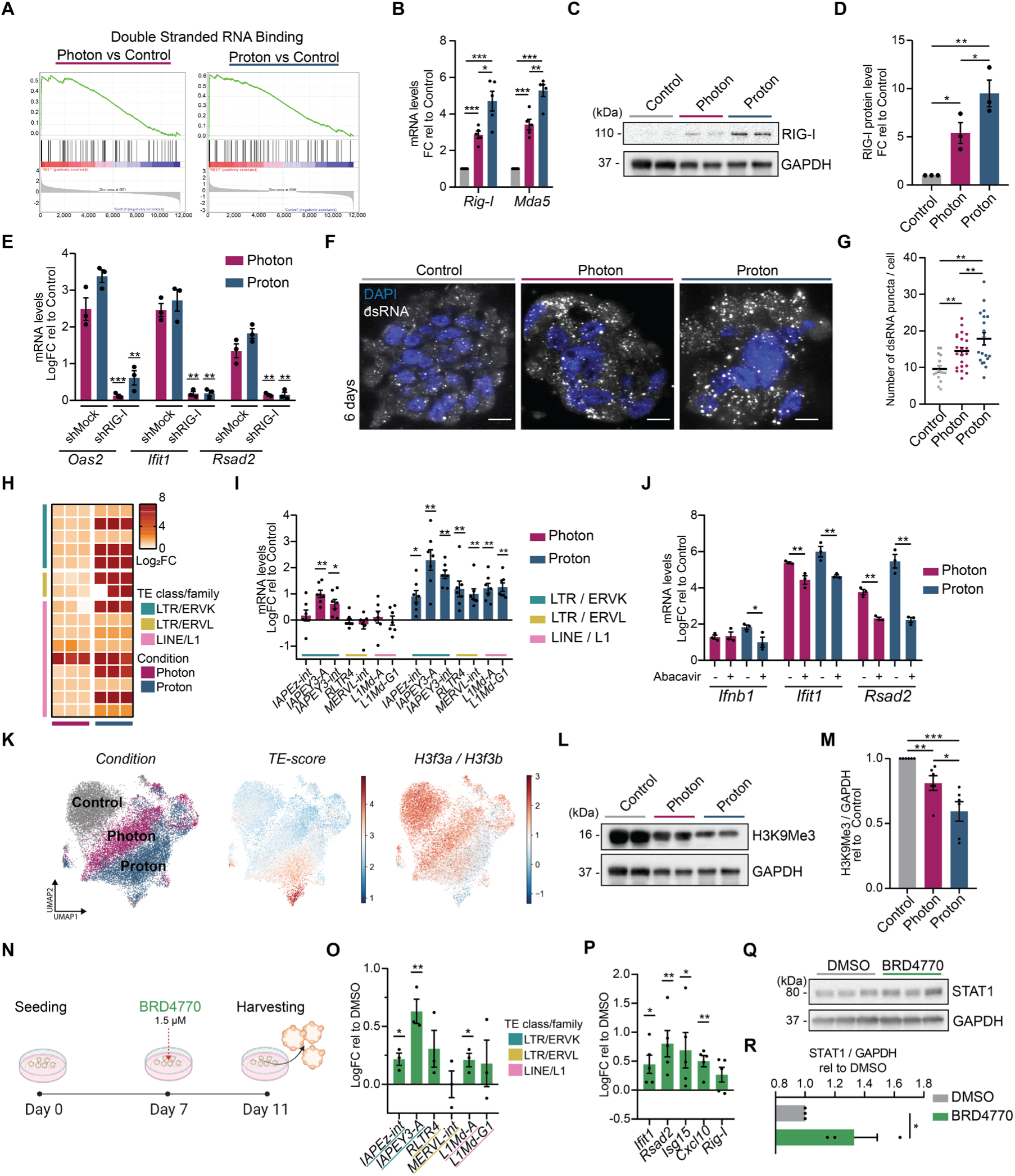
Irradiation leads to dsRNA accumulation into the cytoplasm and TE upregulation. (**A**) Enrichment plots of the biological process double stranded RNA binding at 6 days post photon and proton irradiation. (**B**) rt-qPCR analysis of *Rig-I* and *Mda5* at 6 days post irradiation. Data is shown as FC relative to control (means ± s.e.m; n = 5 animals/condition). One-way ANOVA with Tukey’s multiple comparison test. (**C**) Western blot analysis of RIG-I and GAPDH of organoids at 6 days post irradiation. All biological replicates are shown in fig. S4C. (**D**) Western blot quantification of RIG-I at 6 days post irradiation (Fig. 3C and fig. S4C). Data is shown as FC relative to control (means ± s.e.m; n = 3 animals/condition). One-way ANOVA with Tukey’s multiple comparison test. (**E**) rt-qPCR analysis of ISGs in shMock and shRIG-I-treated organoids at 6 days post irradiation. Data is shown as Log_2_FC relative to shMock control (means ± s.e.m; n = 3 animals/condition). One-way ANOVA with Tukey’s multiple comparison test. **p and ***p are relative to shMock. (**F**) Representative images of immunofluorescence staining of organoids sections at 6 days post irradiation. Scale bar, 10 µm. (**G**) Immunofluorescence staining quantification showing the number of dsRNA puncta (means ± s.e.m; dots represent number of organoids counted/condition from 3 different animals). One-way ANOVA with Tukey’s multiple comparison test. (**H**) Gene expression level of significant (p < 0.05) TEs extrapolated from the bulk RNA-seq data of organoids at 6 days post irradiation. Data is shown as Log_2_FC relative to control samples. (**I**) rt-qPCR analysis of TEs at 6 days post irradiation. Data is shown as a FC relative to control (means ± s.e.m; n = 7 animals/condition). Two-way ANOVA with Tukey’s multiple comparison test. (**J**) rt-qPCR analysis of ISGs in Abacavir treated (+) and untreated (-) organoids at 6 days post irradiation. Data is shown as Log_2_FC relative to Control (means ± s.e.m; n = 3 animals/condition). Two-sided unpaired *t*-test. (**K**) UMAPs of the non-integrated dataset of organoids at 6 days post irradiation (day 11) re-analyzed considering both genes and TEs showing: the different conditions (left panel); cells expressing *MTE2b, B2_Mm2, RSINE1, M1_Mus2, B1_Mus1* and *B3* (TE-score) (central panel); cells expressing *H3f3a* and *H3f3b* (right panel). Individual genes and TEs are shown in fig. S4, G to I. (**L**) Western blot analysis of H3K9Me and GAPDH of organoids at 6 days post irradiation. All biological replicates are shown in fig. S5A. (**M**) Western blot quantification of H3K9Me3 (Fig. 3L and fig. S5A). H3K9Me3 protein levels were normalized for GAPDH (means ± s.e.m; n = 6 animals/condition). One-way ANOVA with Tukey’s multiple comparison test. (**N**) Schematic representation of BRD4770 treatment. Organoids were treated at day 7 and harvested at day 11. (**O**) rt-qPCR analysis of TEs in BRD4770 and DMSO-treated organoids at day 11. Data is shown as Log_2_FC relative to DMSO (means ± s.e.m; n = 3 animals/condition). Two-sided unpaired *t*-test. (**P**) rt-qPCR analysis of ISGs in BRD4770 and DMSO-treated organoids at day 11. Data is shown as Log_2_FC relative to DMSO (means ± s.e.m; n = 5 animals/condition). Two-sided unpaired *t*-test. (**Q**) Western blot analysis of STAT1 and GAPDH in BRD4770 and DMSO treated-organoids at day 11. (**R**) Western blot quantification of STAT1 (Figure 3Q). STAT1 protein levels were normalized for GAPDH (means ± s.e.m; n = 3 animals/condition). Two-sided unpaired *t*-test. In all figures *p < 0.05, **p < 0.01, ***p < 0.005.

Since mitochondrial damage has been linked to inflammation and IFN-I signaling activation (*32*, *33*), and given the low levels of micronuclei observed at later time points (Fig. 2B), we investigated whether mitochondrial dysfunction and leakage of mitochondrial DNA (mtDNA) into the cytoplasm could explain the sustained IFN-I response detected 6 days post-irradiation. Although micronuclei levels remained low at this timepoint (Fig. 2B), dot blot analysis of cytoplasmic fractions revealed persistently high levels of dsDNA after irradiation (fig. S3D). Moreover, the level of cytosolic ZBP1, a cytosolic nucleic acid sensor that modulates the immune response upon recognition of different forms of dsDNA, including mtDNA (*33*, *34*), were significantly increased in irradiated organoids (Fig. 2, G and H and fig. S3E). In line with this, flow cytometry analysis revealed a decrease in mitochondrial membrane potential (mtMP) (Fig. 2I right) and an increase in total mitochondrial mass (Fig. 2I left), indicating the accumulation of dysfunctional mitochondria (*35*). This phenotype was further accompanied by a significant increase in cytosolic mtDNA levels (Fig. 2J). Depletion of mtDNA by treatment with non-cytotoxic concentrations of ethidium bromide (EtBr) (*36*) (fig. S3, F and G) resulted in a pronounced decrease in ISG expression following both photon and proton irradiation (Fig. 2K and fig. S3H). Additionally, ZBP1 levels were reduced in EtBr-treated samples (fig. S3, I and J), confirming a link between mtDNA, ZBP1 and ISG upregulation. Together, these findings demonstrate that cGAS plays an essential role in the regulation of IFN-I signaling after radiation-induced DNA damage in normal tissue cells, while ZBP1 may further reinforce the inflammatory response by sensing mtDNA. However, the activation of these pathways does not explain the differential inflammatory response observed after proton irradiation.

### Radiation-induced loss of histone H3 methylation leads to derepression of transposable elements and cytoplasmic dsRNA formation

Gene set enrichment analysis (GSEA) revealed a significant upregulation of double-stranded (ds)RNA binding components at 6 days after irradiation (Fig. 3A and fig. S4A). Unlike dsDNA sensors, several genes associated with this GO term exhibited a higher expression upon proton compared to photon irradiation (fig. S4B). Among these, the Rig-I like Receptors (RLRs) *Mda5* (or *Ifih1*) and *Rig-I* (or *Ddx58*), key modulators of the inflammatory response induced by detection of cytoplasmic dsRNA (*37*, *38*), were found to be highly upregulated especially with protons (Fig. 3B and fig. S4B). Western blot analysis further confirmed a significantly higher increase of RIG-I at 6 days after 7 Gy proton irradiation (Fig. 3, C and D and fig. S4C). Accordingly, RIG-I knockdown resulted in a marked reduction of ISG expression (Fig. 3E), underscoring the critical role of this cytoplasmic dsRNA sensor in regulating IFN-I signaling after irradiation (*39*). Microscopy analysis of organoid sections revealed a pronounced accumulation of dsRNA in the cytoplasm of irradiated samples (Fig. 3F), supporting the observed upregulation of dsRNA sensors. Notably, 7 Gy proton-irradiated organoids exhibited more and bigger dsRNA foci (Fig. 3G and fig. S4D), reinforcing the link between the increased upregulation of RIG-I and the higher IFN-I response detected upon proton irradiation.

The derepression of TEs, such as endogenous retroviruses (ERVs), has been implicated in endogenous dsRNA accumulation (*40–42*). Indeed, bulk RNA-seq and qPCR analyses revealed a significant upregulation of multiple TEs, including ERVs (such as ERVK and ERVL) and LINE-1 (L1) elements, 6 days after irradiation, with a more pronounced increase following proton irradiation (Fig. 3, H and I). Consistently, inhibiting downstream effectors of ERV activation using the nucleoside reverse transcriptase inhibitor Abacavir (*43*) (fig. S4E), reduced IFN-I signaling (Fig. 3J), thereby supporting the role of epigenetic derepression-mediated ERV activation in IFN-I signaling post-irradiation. The expression of ERV and L1 elements is tightly regulated by several regulatory networks, with histone variant H3.3 proteins (*H3f3a* and *H3f3b*) and their methylation playing a pivotal role in their silencing. Loss of H3.3 or the repressive histone mark H3K9Me have been linked to TE derepression (*41*, *43–45*). In accordance with this, bulk RNA-seq analysis showed a strong downregulation of histone variants *H3f3a* and *H3f3b*, along with several methyltransferase-related genes (fig. S4F). scRNA-seq further corroborated this findings, showing a pronounced increased of several TEs after proton irradiation (Fig. 3K and fig.S4, G and H), including the *B2_Mm2* TE, which was shown to be a potent IFN-β enhancer (*46*). This aligned with a pronounced decrease of *H3f3a* and *H3f3b* expression (Fig. 3K right and S4I). Moreover, western blot analysis revealed a global reduction of H3K9Me upon irradiation, especially with protons (Fig. 3, L and M and fig. S5, A and B). Given H3K9Me’s crucial role in ERV and L1 element regulation, we next treated non-irradiated organoids with the H3K9Me transferase inhibitor BRD4770 (Fig. 3N). BRD4770 treatment led to upregulation of ERVK, ERVL and L1 elements (Fig. 3O), increased ISG expression (Fig. 3P), and elevated STAT1 protein levels (Fig. 3, Q and R), highlighting the role of histone H3 methylation in regulating TE activation and subsequent dsRNA-mediated IFN-I signaling pathway. Taken together, these data indicate the importance of TE deregulation in IFN-I signaling activation and suggest that the heightened response to protons can be attributed to the more pronounced loss of H3K9Me, leading to increased TE-related dsRNA formation and IFN-I response.

### INF-β enhances salivary gland stem/progenitor cell activity after irradiation

Radiation-induced genomic instability and changes in the microenvironment can profoundly affect the self-renewal capacity and longevity of stem cells (*47*, *48*). Given the stronger IFN-I response and self-renewal capacity observed upon proton irradiation, along with the previously observed role of IFN-β in promoting regeneration and stem cell function post-irradiation (*28*), we investigated whether IFN-β in the irradiated salivary gland microenvironment influences the self-renewal potential of salivary gland stem/progenitor cells. Since proton-irradiated organoids already exhibited high IFN-β levels, recombinant IFN-β or IFN-β-neutralizing antibody (IFN-β-NA) were added to the culture media after photon irradiation and the stem/progenitor self-renewal capacity was assessed (Fig. S5C). Interestingly, IFN-β treatment significantly enhanced cell proliferation, as shown by the increased cell number after treatment (Fig. 4, A and B). Additionally, IFN-β blocking decreased organoid formation (Fig. 4, C and D), confirming a role for IFN-β in regulating stem/progenitor cell activity after irradiation.

**Figure 4:**
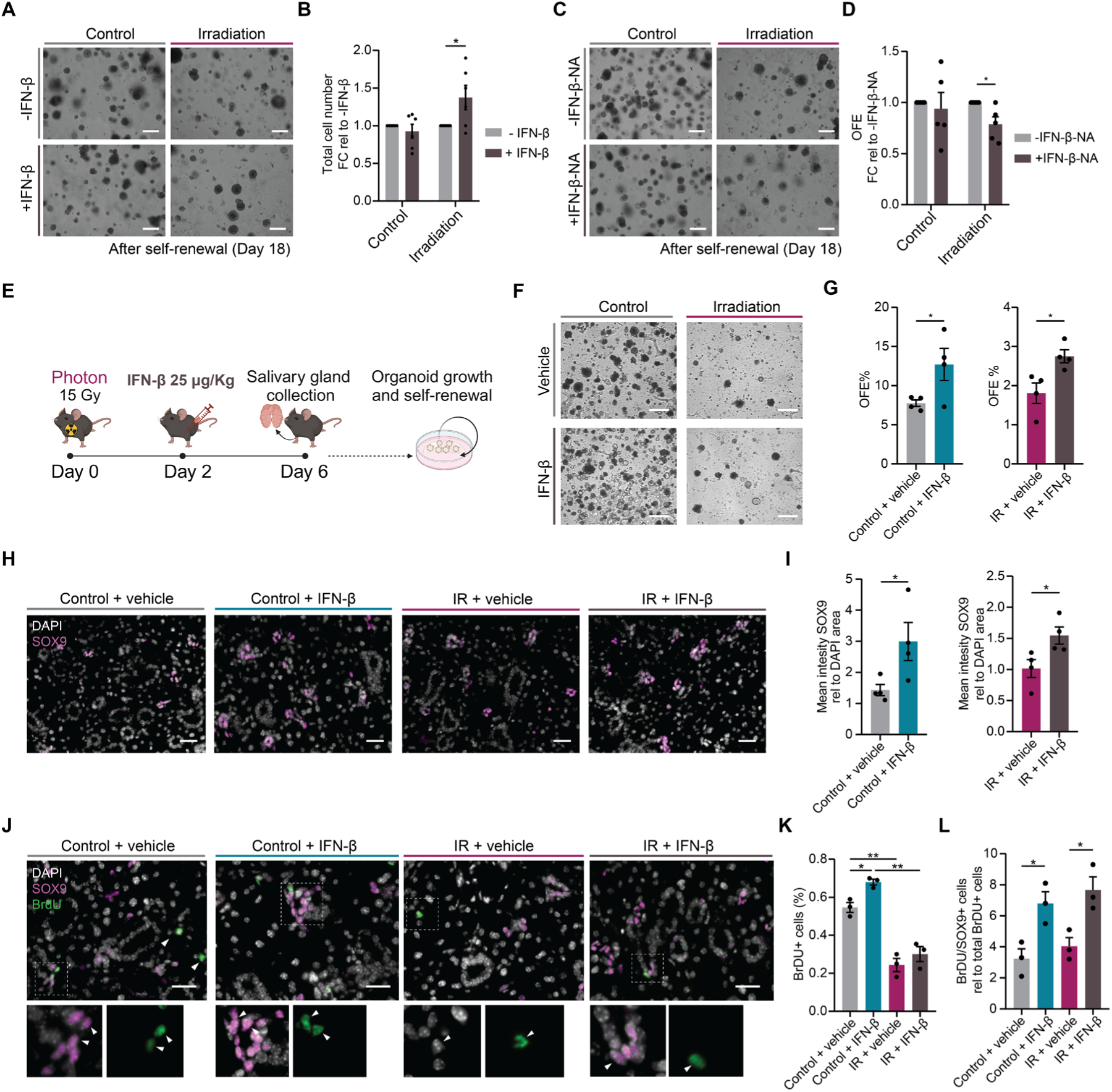
IFN-β promotes stem/progenitor cell activity. (**A**) Representative images of control and irradiated organoids in culture after self-renewal (day 18) and treatment with (+) or without (-) IFN-β. Scale bar, 100 µm. (**B**) Quantification of the total number of cells of control and irradiated organoids after self-renewal (day 18) and treatment with (+) or without (-) IFN-β. Data is shown as FC relative to untreated organoids (means ± s.e.m; n = 6 animals/condition). Two-sided unpaired *t*-test. (**C**) Representative images of control and irradiated organoids in culture after self-renewal (day 18) and treatment with (+) or without (-) IFN-β-NA. Scale bar, 100 µm. (**D**) OFE of control and irradiated organoids after self-renewal (day 18) and treatment with (+) or without (-) IFN-β-NA. Data is shown as FC relative to untreated organoids (means ± s.e.m; n = 5 animals/condition). Two-sided unpaired *t*-test. (**E**) Schematic representation of the animal experiment. (**F**) Representative images of organoids derived from control and irradiated mice treated with vehicle or IFN-β. Scale bar, 100 µm. (**G**) OFE% of organoids derived from control and irradiated mice treated with vehicle or IFN-β (means ± s.e.m; n = 4 animals/condition). Two-sided unpaired *t*-test. (**H**) Representative images of immunofluorescence staining of salivary gland sections from control and irradiated mice treated with vehicle or IFN-β. Scale bar, 25 µm. (**I**) Immunofluorescence staining quantification showing SOX9 mean intensity (means ± s.e.m; n = 4 animals/condition). Two-sided unpaired *t*-test. (**J**) Representative images of immunofluorescence staining of salivary gland sections from control and irradiated mice treated with vehicle or IFN-β. Zoom-ins show SOX9- and BrDU-expressing cells. Scale bar, 25 µm. (**K**) Immunofluorescence staining quantification showing the percentage (%) of BrDU+ cells (means ± s.e.m; n = 4 animals/condition). Two-sided unpaired *t*-test. (**L**) Immunofluorescence staining quantification showing SOX9/BrDU+ cells. Data is relative to total BrDU+ cells (means ± s.e.m; n = 4 animals/condition). Two-sided unpaired *t*-test. In all figures *p < 0.05, **p < 0.01

To validate these findings *in vivo*, we used our established mouse model (*20*, *49*). Mice received a single intraperitoneal dose of IFN-β [25 µg/kg (*28*)] 2 days after local irradiation (IR) with 15 Gy, and salivary glands were collected on day 6 (Fig. 4E). While irradiation caused a slight decrease in body weight, no significant differences were observed in the irradiated salivary glands compared to controls (fig. S5, D and E). Similar to irradiated organoids, cells derived from irradiated salivary glands exhibited reduced OFE; however, IFN-β treatment significantly improved organoid formation compared to vehicle-treated glands (Fig. 4, F and G), confirming its positive effects in stem/progenitor activity.

Previously, we identified a *Sox9*-expressing stem/progenitor cell population with high IFN-I signaling and enhanced proliferation following irradiation (Fig. 1, L and M). Furthermore, scRNA-seq analysis of locally irradiated mice [E-MTAB-1374 (*50*)] revealed an *Epcam+* salivary gland population expressing high *Sox9* levels in both control and irradiated conditions, enriched with proliferation markers such as *Ccnd1* (Cyclin D1) (Fig. S5F), aligning with our *in vitro* findings. Given the role of this population in organoid formation, self-renewal (*18*), and tissue regeneration after injury (*51–53*), we investigated whether functional changes following irradiation and IFN-β treatment could account for the observed differences in organoid formation. Interestingly, IFN-β treatment led to a significant increase of SOX9-expressing cells in both control and irradiated mice (Fig. 4, H and I and fig. S5G). Additionally, although irradiation caused a pronounced decrease in proliferating cells (Fig. 4, J and K), IFN-β-treated mice showed an increase in proliferating SOX9-expressing cells (Fig. 4L), further supporting our previous observations. Together, these data confirm the beneficial effects of IFN-β in maintaining salivary gland stem/progenitor cells and enhancing their proliferative capacity.

## Discussion

In this study, we determined the molecular mechanisms underlying IFN-I signaling activation in normal tissue-derived salivary gland organoids following photon and proton irradiation. Our findings reveal a link between radiation-induced chromatin modification, TE derepression and cytoplasmic dsRNA formation, culminating in the upregulation of ISGs and enhanced stem/progenitor cell activity and self-renewal capacity. Moreover, using bulk and single cell transcriptomics we demonstrated that proton irradiation leads to a more pronounced upregulation of IFN-I signaling, resulting in a stronger activation of *Sox9*-expressing stem/progenitor cells compared to conventional photon irradiation.

Activation of the cGAS-STING pathway following micronuclei formation is a major driver of IFN-I induction upon radiation-induced DNA damage in cancer cells and transformed cell lines (*30*). While a recent study showed that genotoxic stress-induced micronuclei do not activate cGAS in cell lines (*54*), our work using a salivary gland organoid model, reflective of normal tissue stem cells (*8*, *16*, *18*, *55*, *56*), demonstrates a significant increase of cytoplasmic cGAS and micronuclei formation at 2 days after photon and proton irradiation. Knock-down of cGAS abrogates the later IFN-I response, underscoring its essential role in shaping the overall radiation-induced inflammatory response. Interestingly, cGAS levels decrease at 6 days, which can be attributed to the propensity of micronucleated cells to undergo cell death or apoptosis (*57*). Moreover, although the fate of micronuclei remains largely unclear, studies suggest that they may undergo degradation or reincorporation into the nucleus over time (*57*, *58*), offering an alternative explanation for their reduced number at later time points.

Similar to nuclear DNA, the release of mtDNA due to mitochondrial dysregulation can also activate cytoplasmic nucleic acid sensors, leading to inflammation (*32*). Our study reveals that photon and proton irradiation induce a simultaneous accumulation of damaged mitochondria and cytoplasmic dsDNA and mtDNA at 6 days post-irradiation. This aligns with previous work by us (*35*) and others (*59*), showing a correlation between the accumulation of elongated-damaged mitochondria and the upregulation of IFN-β following mtDNA leakage into the cytoplasm. The low levels of cGAS, however, suggest the involvement of other cytoplasmic nucleic acid sensors, such as ZBP1. While a recent study showed a cooperative role of cGAS and ZBP1 in detecting mtDNA in cardiac cells (*33*), our results show that abrogation of ZBP1, along with the depletion of mtDNA, significantly decreases ISG expression upon irradiation. Together this confirms the crucial role of ZBP1 in sustaining the IFN-I response (*33*, *34*) and suggests the existence of a cGAS-independent mechanism. Nevertheless, these results do not explaining the difference in response to protons and photons.

Although the IFN-I response observed after irradiation is largely mediated by the activation of cytoplasmic dsDNA sensors, irradiated organoids displayed increased expression of TEs, accompanied by an accumulation of cytoplasmic dsRNA and upregulation of the dsRNA sensor RIG-I, which were more pronounced after proton irradiation. Elevated levels of TEs, dsRNA, and RIG-I have been shown to enhance the immune response of cancer cells following irradiation (*39*). However, our study reveals that this phenomenon extends beyond tumors, as irradiated non-tumor cells also express these immune-enhancing elements. This emphasizes the pivotal role of TEs in enhancing the IFN-I response not only in tumors and age-associated disorders (*42*, *43*, *60*, *61*) but also in irradiated normal tissue-derived stem/progenitor cells. Importantly, TEs are tightly regulated, and their expression can be profoundly affected by alterations in chromatin accessibility (*41*, *44*, *62*). We demonstrate that proton irradiation leads to a more pronounced loss of histone H3K9Me and downregulation of the histone variants *H3f3a* and *H3f3b* at 6 days, pivotal factors in TE silencing (*63*). Consistent with our findings, previous studies have shown a differential epigenetic modulation and downregulation of histone H3 methylation in various tumors following proton irradiation (*22*).

Although protons elicit a stronger IFN-I response, proton-irradiated organoids show enhanced self-renewal capacity compared to photon-irradiated organoids, reflecting higher stem cell activity and proliferation. Additionally, we show that treatment of photon-irradiated organoids with IFN-β improves their self-renewal, while inhibiting IFN-β signaling impairs their OFE. These findings highlight the crucial role of IFN-β in enhancing stem cell activity, with potential advantages for normal tissue recovery following radiation. Recent studies have demonstrated that the presence of cGAS-STING-dependent-IFN-β in the niche of irradiated-intestinal stem/progenitor cells (*13*), as well as the activation of TEs in chemotherapy-treated hematopoietic stem cells (*27*), improved tissue regeneration. Our scRNA-seq analysis further supports these findings, revealing a cluster of cells with highly upregulated TEs and a robust IFN-I response, which leads to increased proliferation of *Sox9+* stem/progenitor cells following proton irradiation. Additionally, DNA demethylation has been associated with stem cell identity, fate specification and dedifferentiation (*64–66*). However, its precise mechanisms remain unclear. Here, we propose IFN-I as the key intermediate linking radiation-induced damage to epigenetic changes.

While our study establishes a link between higher self-renewal capacity of proton-irradiated organoids and increased upregulation of TEs and related IFN-I response, further studies are warranted to validate the potential clinical implications in an *in vivo* setting and to evaluate the combined effects of IFN-β activation and the immune response. Furthermore, it is also important to assess the effects of alterations to the tumor microenvironment on cancer stem cell proliferation and plasticity. Optimizing the balance between cancer cell elimination and the enhancement of normal stem cell function and tissue regeneration is essential, as it can significantly contribute to improving patient outcome post-cancer treatment.

## Materials and Methods

The details of all antibodies and reagents used in this study are listed in Table S1

### Mice

For the culturing of salivary gland organoids, 8-to 12-week-old female C57BL/6 mice (Envigo, the Netherlands) were used. Mice were housed under conventional conditions with a standard diet and water *ad libitum* at the central animal facility of the University Medical Center Groningen. Animal experimental procedures were approved by the Central Committee of Animal Experimentation of the Dutch government and the Institute Animal Welfare Body of the University Medical Center Groningen [animal welfare body (IVD) protocol number 184824-01-001].

### Mice irradiation and treatments

Salivary glands of 12-week-old female C57BL/6 mice (Envigo, the Netherlands) were (sham-) irradiated with a single dose of 15 Gy X-rays (RAD 320, Precision X-ray) Two days after irradiation, mice were injected with a single intraperitoneal dose of IFN-β (25 µg/kg) (MedChemExpress, HY-P73130) or vehicle (saline solution) and sacrificed 6 days after irradiation. To assess proliferation, all mice were subjected to an intraperitoneal BrdU (Sigma-Aldrich, B5002-1G) injection. BrdU was dissolved in physiological solution at a concentration of 50 mg/kg of body weight and injected 1 day before sacrifice.

### Organoid culture

Following the collection of submandibular salivary glands, tissue isolation was performed as previously described (*49*). In summary, after mechanic and enzymatic digestion, cells were cultured in DMEM/F12 complemented with penicillin-streptomycin (Pen/Strep) antibiotics, glutamax (2 mM), EGF (20 ng/ml); FGF2 (20 ng/ml), N2 (1x), insulin (10 µg/ml) and dexamethasone (1 uM). After 72 hours in culture, primary spheres were dissociated into single cells using 0.05% trypsin EDTA and counted. 10.000 cells were then plated in 75 µL gel/well [35 µL cell suspension + 40 µL of culturex basement membrane extract Type 2 (BME)] or [25 uL cell suspension + 50 uL of Matrigel] in a 12-well tissue culture plate. After solidification of the gels, 1 mL of WRY medium [DMEM/F12, penicillin-streptomycin, glutamax, N2, EGF, FGF2, insulin, Y27632 (10 µM), 10% R-spondin1–conditioned medium, and 50% Wnt3a-conditioned medium] was added to each well. The gels were then incubated at 37°C and 5% CO_2_ until the day of the experiment.

### Organoid irradiation and treatments

*Irradiation:* After 5 days in culture, mouse salivary gland organoids were irradiated with 4, 7 and 15 Gy photons or protons. Photon irradiation was performed using a Cesium-137 source with a dose rate of 0.59 Gy/min at the department of Biomedical Sciences of the UMCG. Proton irradiation was performed as previously described (*67*) using a 150 MeV proton beam at the Particle Therapy Research Center (PARTREC) accelerator facility of the UMCG and at the UMCG Proton Therapy Center (GPTC). Samples were placed in the plateau region of a 150 MeV Bragg curve. To ensure similar culture conditions, photon and proton irradiations were performed on the same day. Plates were sealed at the time of irradiation to ensure sterility and the medium was replaced with fresh medium right after irradiation at day 5. *Drug treatments:* C-176 (0.5 µM) or equivalent volume of vehicle (DMSO) was added to the media of irradiated and non-irradiated organoids at 1 day and 4 days after irradiation; BRD4770 (1.5 µM) or equivalent volume of vehicle (DMSO) was added to the media of 7 day-old non-irradiated organoids; Abacavir (0.2 µM) or equivalent volume of vehicle (PBS) was added to the media of irradiated and non-irradiated organoids at day 6, 8 and 10; IFN-β was added to the media of photon-irradiated and non-irradiated organoids at day 7 at a final concentration of 2 ng/mL and at the time of self-renewal at a final concentration of 0.5 ng/mL; IFN-β-NA was added to the media of photon-irradiated and non-irradiated organoids 1 day after irradiation at a final concentration of 2 µg/mL; and, for an effective depletion of mtDNA, organoids were treated with Ethidium Bromide (EtBr) from day 2 at a final concentration of 100 ng/mL.

### Survival and self-renewal assay after irradiation

Organoids were collected at day 11 (6 days post irradiation) and dissociated into single cells using 0.05% trypsin-EDTA. Organoid number and single cells (trypan blue negative) were noted and used to assess the effects of irradiation on organoid growth and cell survival.

To assess the self-renewal capacity of irradiated-stem/progenitor cells, organoids were harvested at day 11 (6 days post irradiation) and dissociated into single cells using 0.05% trypsin-EDTA. Live cells were counted (trypan blue negative) and 10.000 cells were plated in 75 µL gel/well in a 12-well tissue culture plate with 1 mL of WRY medium. Plates were then incubated at 37°C and 5% CO_2_. One week later (day 18), organoids were harvested and counted. The number of organoids was noted and used to calculate the organoid formation efficiency (OFE) as follows:

*OFE % = (number of organoids harvested / number of cells seeded) x 100*

### Bulk RNA-sequencing library preparation

For the cDNA library preparation, total RNA was extracted from organoids at 2 (day 7) and 6 days (day 11) post irradiation using RNeasy Mini Kit according to the manufacturer’s instructions. RNA quality was determined using High Sensitivity RNA ScreenTapes on an Agilent 2100 Bioanalyzer. Highly intact RNA (RIN value > 9) from organoids derived from 3 different animals/condition was then processed for the cDNA library preparation using the QuantSeq 3’ mRNA-Seq Library Prep Kit following the manufacturer’s protocol; 100 ng of RNA was used as an input for all samples. Libraries were equimolarly pooled and 1.8 pM of the pool with 15% PhiX were loaded on a NextSeq 500 (Illumina) for a 75 bp single-read sequencing run at the Research Sequencing Facility of ERIBA (UMCG).

### Bulk RNA-sequencing analysis

For the alignment of the fastq files the mouse genome GRCm39 was used. The gene counts per sample were detected with the QuantSeq 3’ mRNA-seq Integrated Data Analysis Pipeline on the BlueBee Genomics Platform provided by Lexogen with default settings. R (v4.1.3) and RStudio were used for the downstream analysis and plots were generated with the R package ggplot2 (*68*). For the differential expression analysis, low expressed genes (total count <1 in less than 2 samples) were excluded. The R package edgeR (*69*) was used for normalization and identification of differential expressed genes (LogFC > 0.6, FDR < 0.05). Gene set enrichment analysis (GSEA) for differentially expressed genes and reduction of redundant gene ontology (GO) terms were performed with the R packages clusterProfiler (*70*) and Revigo (*71*). GO terms with an adjusted *P* value < 0.05 were considered significant.

For the identification of TEs, the R package Rsubread (*72*) was used for the annotation upon alignment of the fastq files on the mouse genome GRCm39. The database gEVE version 1.1 was used to retrieve the annotation datasheet. The R package edgeR was then used for downstream analysis and identification of differentially expressed TEs (logFC > 0.6, FDR < 0.05).

### Western blot

After harvesting, organoids were lysed using RIPA buffer, sonicated and spun down at 4°C (for 5 min at max speed). The protein concentration of the lysates was measured using the Bradford quantification method and the samples were boiled at 99°C for 5 min before loading. An equal amount of protein was separated with 10 or 12% polyacrylamide gels. The transfer to nitrocellulose membranes was performed using the Trans-Blot Turbo System (Bio-Rad). Blocking was performed using 10% milk in PBS-Tween20 for 30 min. After cutting, the membranes were incubated with primary antibodies at 4°C overnight followed by incubation with horseradish peroxidase-conjugated secondary antibodies at room temperature for 1.5 hours. Membranes were developed using ECL reagent in a ChemiDoc imager (Bio-Rad). Western blots analysis was performed using Image Lab software. The list of primary antibodies and dilutions are specified in Table S1.

### Quantitative real-time qPCR

Total RNA was extracted from organoids by using RNeasy Mini Kit according to the manufacturer’s instructions. cDNA reverse transcription was performed by using 1 µL dNTP mix (10 mM), 1 µL random primers (100 ng), 4 µL 5x First-strand Buffer, 2 µL DTT (0.1 M), 1 µL RNase OUTTM (40 units/µL), and 1 µL M-MLV RT (200 units). To assess the expression of the genes of interest, specific primers were used together with iQ SYBR Green Supermix. All reactions were run in triplicate on a Bio-Rad Real-Time PCR System. The list of primers is specified in Table S2. The *Ywhaz* gene was used as internal control.

### Immunofluorescence staining

For the staining of micronuclei, 7- and 11-day-old salivary gland organoids were harvested and dissociated into single cells. After counting, cells were re-seeded at a concentration of 90.000 cells/well on the top of a coverslip in a 24 well plate. WRY media was added and cells were incubated at 37°C and 5% CO_2_ overnight. On the next day cells were fixed with 4% formaldehyde and incubated with cGAS primary antibody followed by incubation with the proper secondary antibody, both at room temperature for 1 hour. DAPI was used for nuclear staining. Images were acquired with a Leica DM6 microscope.

For the staining of organoid sections, 7- and 11-day-old organoids were harvested and fixed with 4% formaldehyde. The organoids were then processed for embedding in paraffin and cut into 4 µm thick sections. For the staining of cGAS and dsRNA, sections were dewaxed and incubated with the proper primary antibody at 4°C overnight. On the next day, sections were incubated with the secondary antibody at room temperature for 1 hour. DAPI was used for nuclear staining. For the co-staining of SOX9 and STAT1 the Opal 4-Color anti-Rabbit Manual IHC Kit was used following the manufacturer’ protocol. The Opal520 fluorophore was used to visualize STAT1 while the Opal570 fluorophore was used to visualize H3K9Me. Spectral DAPI was used for nuclear staining. Images were acquired with a Leica SP8X confocal microscope. The list of antibodies and dilutions are specified in Table S1. Images were analyzed using ImageJ (v1.52). Puncta analysis was performed using Icy (v2.4.2.0).

For the staining of salivary gland tissue, glands were collected and fixed with 4% formaldehyde. Fixed salivary glands were then embedded in paraffin and cut into 4 µm thick sections. Staining was performed as previously described (*49*). Briefly, sections were dewaxed, subjected to antigen retrieval in citric acid buffer, and permeabilized with 0.4% Triton. Sections were then incubated with 2.5 M HCl for 20 minutes, followed by 0.1 M Sodium Borate for 5 minutes at room temperature. Sections were then processed as described for organoid sections.

### Subcellular fractionation

After harvesting, organoids were resuspended in subcellular fractionation buffer [Sucrose (250 mM), HEPES pH 7.4 (20 mM), KCl (10 mM), MgCl_2_, EDTA (1 mM), EGTA (1 mM) and DTT (1 µM)], passed through a needle 10 times and placed on ice for 30 min. The lysates were then centrifuged at 720 xg for 5 min at 4°C. The supernatant containing the cytoplasm fraction was collected and spun at 10.000 xg for 5 min at 4°C. The supernatant was then concentrated through an Amicon Ultra-4 Filter Unit (Merck, cat#UFC8100214). The validation of the cytoplasm fraction was performed by western blot using antibodies against markers for nucleus (H2A), mitochondria (TOMM20) and cytoplasm (GAPDH). A whole cell extract from control samples was used as a positive control.

### Mitochondrial DNA content

Total genomic DNA was obtained from the cytosolic fractions of control and irradiated organoids using the Monarch Genomic DNA Purification Kit according to the manufacturer’s protocol. DNA was then amplified using specific oligos for different mitochondrial DNA-related genes. Their mean expression was used to estimate the mtDNA content in cytoplasmic fractions. All reactions were run in triplicate on a Bio-Rad Real-Time PCR System. *Ywhaz* and *Tert* genes were used as internal controls.

### Dot blot

For the qualitative detection of dsDNA, the cytoplasm fractions from all the conditions were normalized based on their protein concentration (measured using the Bradford quantification assay). 2 µL of cytoplasm lysate was then placed on a nitrocellulose membrane and let dry at room temperature. After blocking with 10% milk in PBS-Tween20 at room temperature for 45 min, the membrane was incubated with dsDNA primary antibody for 1 hour followed by incubation with horseradish peroxidase-conjugated secondary antibody for 1 hour at room temperature. Membranes were developed using ECL reagent in a ChemiDoc imager. Dot blot analysis was performed using Image Lab software. The list of primary antibodies and dilutions are specified in Table S1.

### Lentiviral production and transduction

For the lentiviral production HEK293T cells were plated with DMEM supplemented with 10% FBS, Pen/Strep and glutamax. Cells were then transfected with 3 µg of vector of interest (pLV[shRNA]-EGFP:T2A:Puro-U6>[shRNA]) (Vector Builder), 3 µg of packaging construct (PAX2), 0.7 µg of glycoprotein envelope plasmid (VSV-G) and 40 µL of PEI (1 µg/mL). On the next day, the media was replaced with DMEM/F12 and the viral supernatant was collected after 1 day.

For the transduction, salivary gland organoids were released from the gels and dissociated into single cells. After counting, the cell suspension was incubated with the viral supernatant and polybrene (6 µg/mL) on a 24-well plate overnight at 37°C and 5% CO_2_. On the next day, cells were counted and seeded in BME into a 12-well plate with WRY media. 2 days after seeding the media was supplemented with Puromycin (2 µg/mL) for the selection of the transduced cells and refreshed every 2 days. The efficiency of transduction was >95% for all the conditions and was assessed measuring the percentage of GFP positive cells.

### Flow cytometry

To assess the total amount of mitochondria and their function following photon and proton irradiation, the dyes TMRE and Mitotracker Green were used. Briefly, salivary gland organoids were harvested, dissociated into single cells and resuspended with fresh culture medium supplemented with the probes at a final concentration of 200 nM. Cells were then incubated at 37°C and 5% CO_2_ for 30 min and washed with PBS/BSA 0.2%. Cells were resuspended in PBS/BSA 0.2% and analyzed using the Quanteon flow cytometer at the Flow Cytometry Unit of the UMCG. Data was analyzed using the FlowJo software (v10.7.2).

### Single cell RNA-sequencing library preparation

The preparation of the single cell RNA-sequencing library was performed by using the 10X Chromium Next GEM Single Cell 3’ Kit v3.1, the 10X Chromium Next GEM Chip G Single Cell Kit and the 10X Single Index kit T Set A following the manufacturer’s instructions. In brief, salivary gland organoids were collected and dissociated into single cells. On the same day, for the exclusion of death cells, 45.000 propidium iodide (PI) negative cells/samples were sorted using the Moflo Astrios sorter at the Flow Cytometry Unit at the UMCG. Organoids derived from 4 different mice/conditions were used and pooled at the time of the sequencing. After sorting, samples were loaded on the Chromium Next GEM Chip G and ran in the Chromium Controller (10X Genomics). On the next day, cDNA Amplification and library construction was performed following the manufacturer’s protocol (10X Genomics). Libraries quality control and concentration measurements were performed using the Qubit and Tapestation machines at the Research Sequencing Facility of ERIBA at the UMCG. For the sequencing, libraries were equimolarly pooled and 1.8 pM of the pool with 5% PhiX were loaded on a NextSeq 500 (Illumina) for a 75 bp paired-end sequencing run at the Research Sequencing Facility of ERIBA (UMCG).

### Single cell RNA-sequencing analysis

Reads without cell barcodes or UMIs were excluded and remaining raw reads were aligned to the mouse genome GRCm39. Following demultiplexing and cell filtering, around 45.000 cells were used for downstream analysis (control 14.266 cells; photons 15.926 cells; protons 14.525 cells). Single cell RNA-sequencing analysis was done on R (v4.1.3) using the Seurat V4 (*73*) package. Cells with more than 5% of mitochondria counts and less than 2500 feature counts were filtered out. Elbow plots and Jack Straw were used for the determination of the principal components (PC) to use for UMAP dimensionality reduction. Clusters with less than 100 cells and no significant markers were excluded from the analysis and cluster-markers were identified using the FindAllMarkers function with the statistical test MAST.

Alignment and detection of TEs was performed on Python (v3.0) using the package scTE (*74*). The downstream analysis was then performed using the Scanpy (*75*) package considering both genes and TEs. Clusters and marker genes/TEs were determined using the Leiden algorithm and the Wilcoxon statistical test.

### Statistical analysis

The number of biological replicates and the details of the statistical analyses performed for each experiment are stated in the main and supplementary figure legends. Data is shown as mean ± s.e.m. The Shapiro-Wilk test was used on raw data to test normality of distribution. For the comparison of two groups, two-sided unpaired *t*-test was used. For multiple group comparison, one-way ANOVA and two-way ANOVA with post-hoc Tukey’s test were used. Sample sizes were estimated empirically and p-values ≤ 0.05 were considered statistically significant. In each experiment the measurements were taken from different biological replicates and no samples were measured repeatedly. Statistical analyses were performed using the GraphPad Prism software (v8.0.1).

## Supplementary Materials

Fig. S1 to S5

Table S1 to S2

References (1)

## Funding

This work was supported by the Dutch Cancer Society KWF Grant nr 12092.

## Author contributions

DC, LB and RPC conceived and designed the study. DC, RvdW, MB, ASG and ALJdB performed the experiments and analyzed the data. DC and MJvG performed the irradiations. DC and UB performed the animal experiments. DC, LB and RPC wrote the manuscript with advice and input from MATMvV. All the authors approved the manuscript.

## Competing interests

The authors declare no competing interests.

## Data and materials availability

The bulk RNA-seq and scRNA-seq data have been deposited in the Gene Expression Omnibus (GEO) repository; GSE numbers are available for review purposes and will be made fully available upon acceptance of the manuscript. All other data are stored at the department of Biomedical Sciences, UMCG and are available from the corresponding authors upon request. The details of all antibodies and reagents used in this study are listed in Table S1.

## Notes

### Competing Interest Statement

The authors have declared no competing interest.

### Summary of Updates

Figures, authors and result section have been updated

## References

1. . D. Cinat, R. P. Coppes, L. Barazzuol, DNA Damage-Induced Inflammatory Microenvironment and Adult Stem Cell Response. Front. Cell Dev. Biol. 9, 729136 (2021).

2. J. J. Bednarski, B. P. Sleckman, At the intersection of DNA damage and immune responses. Nat. Rev. Immunol. 19, 231–242 (2019).

3. M. McLaughlin, E. C. Patin, M. Pedersen, A. Wilkins, M. T. Dillon, A. A. Melcher, K. J. Harrington, Inflammatory microenvironment remodelling by tumour cells after radiotherapy. Nature Research (2020). 10.1038/s41568-020-0246-1.

4. V. Klapp, B. Álvarez-Abril, G. Leuzzi, G. Kroemer, A. Ciccia, L. Galluzzi, The DNA Damage Response and Inflammation in Cancer. Cancer Discov. 13, 1521–1545 (2023).

5. J. E. Leeman, P. B. Romesser, Y. Zhou, S. McBride, N. Riaz, E. Sherman, M. A. Cohen, O. Cahlon, N. Lee, Proton therapy for head and neck cancer: expanding the therapeutic window. Lancet Oncol. 18, e254–e265 (2017).

6. P. Dirix, S. Nuyts, W. Van Den Bogaert, Radiation-induced xerostomia in patients with head and neck cancer: A literature review. Cancer 107, 2525–2534 (2006).

7. N. Li, H. Clevers, Coexistence of quiescent and active adult stem cells in mammals. Science (80-. ). 327, 542–545 (2010).

8. P. Van Luijk, S. Pringle, J. O. Deasy, V. V. Moiseenko, H. Faber, A. Hovan, M. Baanstra, H. P. Van Der Laan, R. G. J. Kierkels, A. Van Der Schaaf, M. J. Witjes, J. M. Schippers, S. Brandenburg, J. A. Langendijk, J. Wu, R. P. Coppes, Sparing the region of the salivary gland containing stem cells preserves saliva production after radiotherapy for head and neck cancer. Sci. Transl. Med. 7 (2015).

9. M. I. van Rijn-Dekker, S. la Bastide-van Gemert, M. A. Stokman, A. Vissink, R. P. Coppes, J. A. Langendijk, P. van Luijk, R. J. H. M. Steenbakkers, Radiation-induced Xerostomia is Related to Stem Cell Dose-dependent Reduction of Saliva Production. Int. J. Radiat. Oncol. Biol. Phys. 000 (2024).

10. A. J. S. Rundberg Nilsson, H. Xian, S. Shalapour, J. Cammenga, M. Karin, IRF1 regulates self-renewal and stress responsiveness to support hematopoietic stem cell maintenance. Sci. Adv. 9, eadg5391 (2023).

11. I. Vitale, G. Manic, R. De Maria, G. Kroemer, L. Galluzzi, DNA Damage in Stem Cells. Mol. Cell 66, 306–319 (2017).

12. Q. Yu, Y. V. Katlinskaya, C. J. Carbone, B. Zhao, K. V. Katlinski, H. Zheng, M. Guha, N. Li, Q. Chen, T. Yang, C. J. Lengner, R. A. Greenberg, F. B. Johnson, S. Y. Fuchs, DNA-Damage-Induced Type I Interferon Promotes Senescence and Inhibits Stem Cell Function. Cell Rep. 11, 785–797 (2015).

13. B. J. Leibowitz, G. Zhao, L. Wei, H. Ruan, M. Epperly, L. Chen, X. Lu, J. S. Greenberger, L. Zhang, J. Yu, Interferon b drives intestinal regeneration after radiation. Sci. Adv. 7, 1–13 (2021).

14. C. Rocchi, L. Barazzuol, R. P. Coppes, The evolving definition of salivary gland stem cells. *npj Regen*. Med. 6, 1–8 (2021).

15. L. S. Y. Nanduri, M. Baanstra, H. Faber, C. Rocchi, E. Zwart, G. De Haan, R. Van Os, R. P. Coppes, Purification and Ex vivo expansion of fully functional salivary gland stem cells. Stem Cell Reports 3, 957–964 (2014).

16. M. Maimets, C. Rocchi, R. Bron, S. Pringle, J. Kuipers, B. N. G. Giepmans, R. G. J. Vries, H. Clevers, G. De Haan, R. Van Os, R. P. Coppes, Stem Cell Reports Ar ticle Long-Term In Vitro Expansion of Salivary Gland Stem Cells Driven by Wnt Signals. doi: 10.1016/j.stemcr.2015.11.009 (2016).

17. I. Orhon, C. Rocchi, B. Villarejo-Zori, P. Serrano Martinez, M. Baanstra, U. Brouwer, P. Boya, R. Coppes, F. Reggiori, Autophagy induction during stem cell activation plays a key role in salivary gland self-renewal. Autophagy 18, 293–308 (2022).

18. D. Cinat, R. Maturi, J. P. Gunawan, A. L. J. Bruin, Notch Signaling Drives Pro-Regenerative and Migratory Traits in Glandular Stem / Progenitor cells. (2024).

19. Y. J. Yoon, D. Kim, K. Y. Tak, S. Hwang, J. Kim, N. S. Sim, J. M. Cho, D. Choi, Y. Ji, J. K. Hur, H. Kim, J. E. Park, J. Y. Lim, Salivary gland organoid culture maintains distinct glandular properties of murine and human major salivary glands. Nat. Commun. 13, 1–16 (2022).

20. X. Peng, Y. Wu, U. Brouwer, T. van Vliet, B. Wang, M. Demaria, L. Barazzuol, R. P. Coppes, Cellular senescence contributes to radiation-induced hyposalivation by affecting the stem/progenitor cell niche. Cell Death Dis. 11, 1–11 (2020).

21. R. Alan Mitteer, Y. Wang, J. Shah, S. Gordon, M. Fager, P. P. Butter, H. Jun Kim, C. Guardiola-Salmeron, A. Carabe-Fernandez, Y. Fan, Proton beam radiation induces DNA damage and cell apoptosis in glioma stem cells through reactive oxygen species. Sci. Rep. 5, 1–12 (2015).

22. I. Schniewind, W. W. Hadiwikarta, J. Grajek, J. Poleszczuk, S. Richter, M. Peitzsch, J. Müller, D. Klusa, E. Beyreuther, S. Löck, A. Lühr, S. Frosch, C. Groeben, U. Sommer, M. Krause, A. Dubrovska, C. von Neubeck, I. Kurth, C. Peitzsch, Cellular plasticity upon proton irradiation determines tumor cell radiosensitivity. Cell Rep. 38 (2022).

23. S. Sioen, O. Vanhove, B. Vanderstraeten, C. De Wagter, M. Engelbrecht, C. Vandevoorde, E. De Kock, M. J. Van Goethem, A. Vral, A. Baeyens, Impact of proton therapy on the DNA damage induction and repair in hematopoietic stem and progenitor cells. Sci. Rep. 13, 1–8 (2023).

24. C. M. Frame, Y. Chen, J. Gagnon, Y. Yuan, T. Ma, A. Dritschilo, D. Pang, Proton induced DNA double strand breaks at the Bragg peak: Evidence of enhanced LET effect. Front. Oncol. 12, 1–8 (2022).

25. D. Carvajal Ibañez, M. Skabkin, J. Hooli, S. Cerrizuela, M. Göpferich, A. Jolly, K. Volk, M. Zumwinkel, M. Bertolini, G. Figlia, T. Höfer, G. Kramer, S. Anders, A. A. Teleman, A. Marciniak-Czochra, A. Martin-Villalba, Interferon regulates neural stem cell function at all ages by orchestrating mTOR and cell cycle. EMBO Mol. Med. 15 (2023).

6. M. A. Florez, K. A. Matatall, Y. Jeong, L. Ortinau, P. W. Shafer, A. M. Lynch, R. Jaksik, M. Kimmel, D. Park, K. Y. King, Interferon Gamma Mediates Hematopoietic Stem Cell Activation and Niche Relocalization through BST2. Cell Rep. 33, 108530 (2020).

7. T. Clapes, A. Polyzou, P. Prater, Sagar, A. Morales-Hernández, M. G. Ferrarini, N. Kehrer, S. Lefkopoulos, V. Bergo, B. Hummel, N. Obier, D. Maticzka, A. Bridgeman, J. S. Herman, I. Ilik, L. Klaeylé, J. Rehwinkel, S. McKinney-Freeman, R. Backofen, A. Akhtar, N. Cabezas-Wallscheid, R. Sawarkar, R. Rebollo, D. Grün, E. Trompouki, Chemotherapy-induced transposable elements activate MDA5 to enhance haematopoietic regeneration. Nat. Cell Biol. 23, 704–717 (2021).

28. L. B. J. Leibowitz, G. Zhao, L. Wei, H. Ruan, M. Epperly, L. Chen, X. Lu, J. S. Greenberger, Zhang, J. Yu, Interferon b drives intestinal regeneration after radiation. Sci. Adv. 7 (2021).

29. M. Maimets, C. Rocchi, R. Bron, S. Pringle, J. Kuipers, B. N. G. Giepmans, R. G. J. Vries, H. Clevers, G. de Haan, R. van Os, R. P. Coppes, Long-Term In Vitro Expansion of Salivary Gland Stem Cells Driven by Wnt Signals. Stem cell reports 6, 150–162 (2016).

30. A. Decout, J. D. Katz, S. Venkatraman, A. Ablasser, The cGAS–STING pathway as a therapeutic target in inflammatory diseases. Nat. Rev. Immunol. 21, 548–569 (2021).

31. J. Nassour, L. Gutierrez Aguiar, A. Correia, T. T. Schmidt, L. Mainz, S. Przetocka, C. Haggblom, N. Tadepalle, A. Williams, M. N. Shokhirev, S. C. Akincilar, V. Tergaonkar, G. S. Shadel, J. Karlseder, Telomere-to-mitochondria signalling by ZBP1 mediates replicative crisis. Nature 614, 767 (2023).

32. S. Marchi, E. Guilbaud, S. W. G. Tait, T. Yamazaki, L. Galluzzi, Mitochondrial control of inflammation. Nat. Rev. Immunol. 23, 159–173 (2023).

33. Y. Lei, J. J. VanPortfliet, Y. F. Chen, J. D. Bryant, Y. Li, D. Fails, S. Torres-Odio, K. B. Ragan, J. Deng, A. Mohan, B. Wang, O. N. Brahms, S. D. Yates, M. Spencer, C. W. Tong, M. W. Bosenberg, L. C. West, G. S. Shadel, T. E. Shutt, J. W. Upton, P. Li, A. P. West, Cooperative sensing of mitochondrial DNA by ZBP1 and cGAS promotes cardiotoxicity. Cell 186, 3013–3032.e22 (2023).

34. B. Szczesny, M. Marcatti, A. Ahmad, M. Montalbano, A. Brunyánszki, S. I. Bibli, A. Papapetropoulos, C. Szabo, Mitochondrial DNA damage and subsequent activation of Z-DNA binding protein 1 links oxidative stress to inflammation in epithelial cells. Sci. Rep. 8, 1–11 (2018).

35. D. Cinat, A. L. De Souza, A. Soto-gamez, A. L. Jellema-de, R. P. Coppes, L. Barazzuol, Mitophagy induction improves salivary gland stem/progenitor cell function by reducing senescence after irradiation. Radiother. Oncol., 110028 (2023).

36. W. Lee, H. I. Choi, M. J. Kim, S. Y. Park, Depletion of mitochondrial DNA up-regulates the expression of MDR1 gene via an increase in mRNA stability. Exp. Mol. Med. 40, 109– 117 (2008).

37. D. T. Thoresen, D. Galls, B. Götte, W. Wang, A. M. Pyle, A rapid RIG-I signaling relay mediates efficient antiviral response. Mol. Cell 83, 90–104.e4 (2023).

38. J. Rehwinkel, M. U. Gack, RIG-I-like receptors: their regulation and roles in RNA sensing. Nature Research (2020). 10.1038/s41577-020-0288-3.

39. J. Du, S. I. Kageyama, R. Yamashita, K. Tanaka, M. Okumura, A. Motegi, H. Hojo, M. Nakamura, H. Hirata, H. Sunakawa, D. Kotani, T. Yano, T. Kojima, Y. Hamaya, M. Kojima, Y. Nakamura, A. Suzuki, Y. Suzuki, K. Tsuchihara, T. Akimoto, Transposable elements potentiate radiotherapy-induced cellular immune reactions via RIG-I-mediated virus-sensing pathways. *Commun*. Biol. 6, 1–13 (2023).

40. Y. G. Chen, S. Hur, Cellular origins of dsRNA, their recognition and consequences. Nat. Rev. Mol. Cell Biol. 23, 286–301 (2022).

41. S. Zhao, J. Lu, B. Pan, H. Fan, S. D. Byrum, C. Xu, A. Kim, Y. Guo, K. L. Kanchi, W. Gong, T. Sun, A. J. Storey, N. T. Burkholder, S. G. Mackintosh, P. C. Kuhlers, R. D. Edmondson, B. D. Strahl, Y. Diao, A. J. Tackett, J. R. Raab, L. Cai, J. Song, G. G. Wang, TNRC18 engages H3K9me3 to mediate silencing of endogenous retrotransposons. Nature 623, 633–642 (2023).

42. K. B. Chiappinelli, P. L. Strissel, A. Desrichard, H. Li, C. Henke, B. Akman, A. Hein, N. S. Rote, L. M. Cope, A. Snyder, V. Makarov, S. Buhu, D. J. Slamon, J. D. Wolchok, D. M. Pardoll, M. W. Beckmann, C. A. Zahnow, T. Mergoub, T. A. Chan, S. B. Baylin, R. Strick, Inhibiting DNA Methylation Causes an Interferon Response in Cancer via dsRNA Including Endogenous Retroviruses. Cell 162, 974–986 (2015).

43. X. Liu, Z. Liu, Z. Wu, J. Ren, Y. Fan, L. Sun, G. Cao, Y. Niu, B. Zhang, Q. Ji, X. Jiang, C. Wang, Q. Wang, Z. Ji, L. Li, C. R. Esteban, K. Yan, W. Li, Y. Cai, S. Wang, A. Zheng, Y. E. Zhang, S. Tan, Y. Cai, M. Song, F. Lu, F. Tang, W. Ji, Q. Zhou, J. C. I. Belmonte, W. Zhang, J. Qu, G. H. Liu, Resurrection of endogenous retroviruses during aging reinforces senescence. Cell 186, 287–304.e26 (2023).

44. N. Dopkins, M. M. O’Mara, E. Lawrence, T. Fei, S. Sandoval-Motta, D. F. Nixon, M. L. Bendall, A field guide to endogenous retrovirus regulatory networks. Mol. Cell 82, 3763– 3768 (2022).

45. S. J. Elsässer, K. M. Noh, N. Diaz, C. D. Allis, L. A. Banaszynski, Histone H3.3 is required for endogenous retroviral element silencing in embryonic stem cells. Nature 522, 240–244 (2015).

46. I. Horton, C. J. Kelly, A. Dziulko, D. M. Simpson, E. B. Chuong, Mouse B2 SINE elements function as IFN-inducible enhancers. Elife 12, 1–24 (2023).

7. F. Lazure, R. Farouni, K. Sahinyan, D. M. Blackburn, A. Hernández-Corchado, G. Perron, T. Lu, A. Osakwe, J. Ragoussis, C. Crist, T. J. Perkins, A. Jahani-Asl, H. S. Najafabadi, V. D. Soleimani, Transcriptional reprogramming of skeletal muscle stem cells by the niche environment. Nat. Commun. 14 (2023).

8. A. J. Wagers, The stem cell niche in regenerative medicine. Cell Stem Cell 10, 362–369 (2012).

9. C. Rocchi, D. Cinat, P. S. Martinez, A. L. Jellema-De Bruin, M. Baanstra, U. Brouwer, C. del Angel Zuivre, H. Schepers, R. van Os, L. Barazzuol, R. P. Coppes, The Hippo signaling pathway effector YAP promotes salivary gland regeneration after injury. Sci. Signal. 14 (2021).

50. N. J. G. McKendrick, G. R. Jones, S. S. Elder, E. Watson, W. T’Jonck, E. Mercer, M. S. Magalhaes, C. Rocchi, L. M. Hegarty, A. L. Johnson, C. Schneider, B. Becher, C. Pridans, Mabbott, Z. Liu, F. Ginhoux, M. Bajenoff, R. Gentek, C. C. Bain, E. Emmerson, CSF1R-dependent macrophages in the salivary gland are essential for epithelial regeneration after radiation-induced injury. Sci. Immunol. 8 (2023).

51. X. Xu, G. Xiong, M. Zhang, J. Xie, S. Chen, K. Li, J. Li, Y. Bao, C. Wang, D. Chen, Sox9+ cells are required for salivary gland regeneration after radiation damage via the Wnt/β-catenin pathway. J. Genet. Genomics 49, 230–239 (2022).

52. S. Chen, K. Li, X. Zhong, G. Wang, X. Wang, M. Cheng, J. Chen, Z. Chen, J. Chen, C. Zhang, G. Xiong, X. Xu, D. Chen, H. Li, L. Peng, Sox9-expressing cells promote regeneration after radiation-induced lung injury via the PI3K/AKT pathway. Stem Cell Res. Ther. 12, 1–13 (2021).

53. S. Aggarwal, Z. Wang, D. R. F. Pacheco, A. Rinaldi, A. Rajewski, J. Callemeyn, E. Van Loon, B. Lamarthée, A. E. Covarrubias, J. Hou, M. Yamashita, H. Akiyama, S. A. Karumanchi, C. N. Svendsen, P. W. Noble, S. C. Jordan, J. J. Breunig, M. Naesens, P. E. Cippà, S. Kumar, SOX9 switch links regeneration to fibrosis at the single-cell level in mammalian kidneys. Science (80-. ). 383, 1–14 (2024).

54. T. Takaki, R. Millar, C. T. Hiley, S. J. Boulton, Micronuclei induced by radiation, replication stress, or chromosome segregation errors do not activate cGAS-STING. Mol. Cell 84, 2203–2213.e5 (2024).

55. R. J. H. M. Steenbakkers, M. I. van Rijn–Dekker, M. A. Stokman, R. G. J. Kierkels, A. van der Schaaf, J. G. M. van den Hoek, H. P. Bijl, M. C. A. Kramer, R. P. Coppes, J. A. Langendijk, P. van Luijk, Parotid Gland Stem Cell Sparing Radiation Therapy for Patients With Head and Neck Cancer: A Double-Blind Randomized Controlled Trial. Int. J. Radiat. Oncol. Biol. Phys. 112, 306–316 (2022).

56. P. W. Nagle, N. A. Hosper, L. Barazzuol, A. L. Jellema, M. Baanstra, M. J. Van Goethem, S. Brandenburg, U. Giesen, J. A. Langendijk, P. Van Luijk, R. P. Coppes, Lack of DNA damage response at low radiation doses in adult stem cells contributes to organ dysfunction. Clin. Cancer Res. 24, 6583–6593 (2018).

57. H. Hintzsche, U. Hemmann, A. Poth, D. Utesch, J. Lott, H. Stopper, Fate of micronuclei and micronucleated cells. Mutat. Res. - Rev. Mutat. Res. 771, 85–98 (2017).

58. M. Kwon, M. L. Leibowitz, J.-H. Lee, Small but mighty: the causes and consequences of micronucleus rupture. Exp. Mol. Med. 52, 1234567890 (2020).

59. A. Irazoki, I. Gordaliza-Alaguero, E. Frank, N. N. Giakoumakis, J. Seco, M. Palacín, A. Gumà, L. Sylow, D. Sebastián, A. Zorzano, Disruption of mitochondrial dynamics triggers muscle inflammation through interorganellar contacts and mitochondrial DNA mislocation. Nat. Commun. 14 (2023).

60. Y. Kong, C. M. Rose, A. A. Cass, A. G. Williams, M. Darwish, S. Lianoglou, P. M. Haverty, A. J. Tong, C. Blanchette, M. L. Albert, I. Mellman, R. Bourgon, J. Greally, S. Jhunjhunwala, H. Chen-Harris, Transposable element expression in tumors is associated with immune infiltration and increased antigenicity. Nat. Commun. 10 (2019).

61. M. De Cecco, T. Ito, A. P. Petrashen, A. E. Elias, N. J. Skvir, S. W. Criscione, A. Caligiana, G. Brocculi, E. M. Adney, J. D. Boeke, O. Le, C. Beauséjour, J. Ambati, K. Ambati, M. Simon, A. Seluanov, V. Gorbunova, P. E. Slagboom, S. L. Helfand, N. Neretti, J. M. Sedivy, L1 drives IFN in senescent cells and promotes age-associated inflammation. Nature 566, 73–78 (2019).

62. T. Kanholm, U. Rentia, M. Hadley, J. A. Karlow, O. L. Cox, N. Diab, M. L. Bendall, T. Dawson, J. I. McDonald, W. Xie, K. A. Crandall, K. H. Burns, S. B. Baylin, H. Easwaran, K. B. Chiappinelli, Oncogenic Transformation Drives DNA Methylation Loss and Transcriptional Activation at Transposable Element Loci. Cancer Res. 83, 2584–2599 (2023).

63. H. A. Lawson, Y. Liang, T. Wang, Transposable elements in mammalian chromatin organization. Nat. Rev. Genet. 24, 712–723 (2023).

64. I. C. MacArthur, L. Ma, C. Y. Huang, H. Bhavsar, M. Suzuki, M. M. Dawlaty, Developmental DNA demethylation is a determinant of neural stem cell identity and gliogenic competence. Sci. Adv. 10, 1–21 (2024).

65. L. Ma, Q. Tang, X. Gao, J. Lee, R. Lei, M. Suzuki, D. Zheng, K. Ito, P. S. Frenette, M. M. Dawlaty, Tet-mediated DNA demethylation regulates specification of hematopoietic stem and progenitor cells during mammalian embryogenesis. Sci. Adv. 8, 1–18 (2022).

66. M. Schulz, A. Teissandier, E. De La Mata Santaella, M. Armand, J. Iranzo, F. El Marjou, P. Gestraud, M. Walter, S. Kinston, B. Göttgens, M. V. C. Greenberg, D. Bourc’his, DNA methylation restricts coordinated germline and neural fates in embryonic stem cell differentiation. Nat. Struct. Mol. Biol. 31, 102–114 (2024).

67. P. W. Nagle, M. J. van Goethem, M. Kempers, H. Kiewit, A. Knopf, J. A. Langendijk, S. Brandenburg, P. van Luijk, R. P. Coppes, In vitro biological response of cancer and normal tissue cells to proton irradiation not affected by an added magnetic field. Radiother. Oncol. 137, 125–129 (2019).

68. H. Wickham, Ggplot2: Elegant Graphics for Data Analysis. Springer-Verlag New York (2016; http://link.springer.com/10.1007/978-0-387-98141-3)vol. 35.

69. M. D. Robinson, D. J. McCarthy, G. K. Smyth, edgeR: A Bioconductor package for differential expression analysis of digital gene expression data. Bioinformatics 26, 139–140 (2009).

70. T. Wu, E. Hu, S. Xu, M. Chen, P. Guo, Z. Dai, T. Feng, L. Zhou, W. Tang, L. Zhan, X. Fu, S. Liu, X. Bo, G. Yu, clusterProfiler 4.0: A universal enrichment tool for interpreting omics data. Innovation 2, 100141 (2021).

71. F. Supek, M. Bošnjak, N. Škunca, T. Šmuc, Revigo summarizes and visualizes long lists of gene ontology terms. PLoS One 6 (2011).

72. Y. Liao, G. K. Smyth, W. Shi, The R package Rsubread is easier, faster, cheaper and better for alignment and quantification of RNA sequencing reads. Nucleic Acids Res. 47 (2019).

73. Y. Hao, S. Hao, E. Andersen-Nissen, W. M. Mauck, S. Zheng, A. Butler, M. J. Lee, A. J. Wilk, C. Darby, M. Zager, P. Hoffman, M. Stoeckius, E. Papalexi, E. P. Mimitou, J. Jain, A. Srivastava, T. Stuart, L. M. Fleming, B. Yeung, A. J. Rogers, J. M. McElrath, C. A. Blish, R. Gottardo, P. Smibert, R. Satija, Integrated analysis of multimodal single-cell data. Cell 184, 3573–3587.e29 (2021).

74. J. He, I. A. Babarinde, L. Sun, S. Xu, R. Chen, J. Shi, Y. Wei, Y. Li, G. Ma, Q. Zhuang, A. P. Hutchins, J. Chen, Identifying transposable element expression dynamics and heterogeneity during development at the single-cell level with a processing pipeline scTE. Nat. Commun. 12, 1–14 (2021).

75. 2* F. Alexander Wolf1*, Philipp Angerer1 and Fabian J. Theis1, SCANPY: large-scale single-cell gene expression data analysis. Genome Biol. 19 (2018).

