## Supplementary Figures and Tables for "Transposable Element Derepression-mediated IFN-I Signaling Enhances Stem/Progenitor Cell Activity After Irradiation"

Cinat D. *et al.*

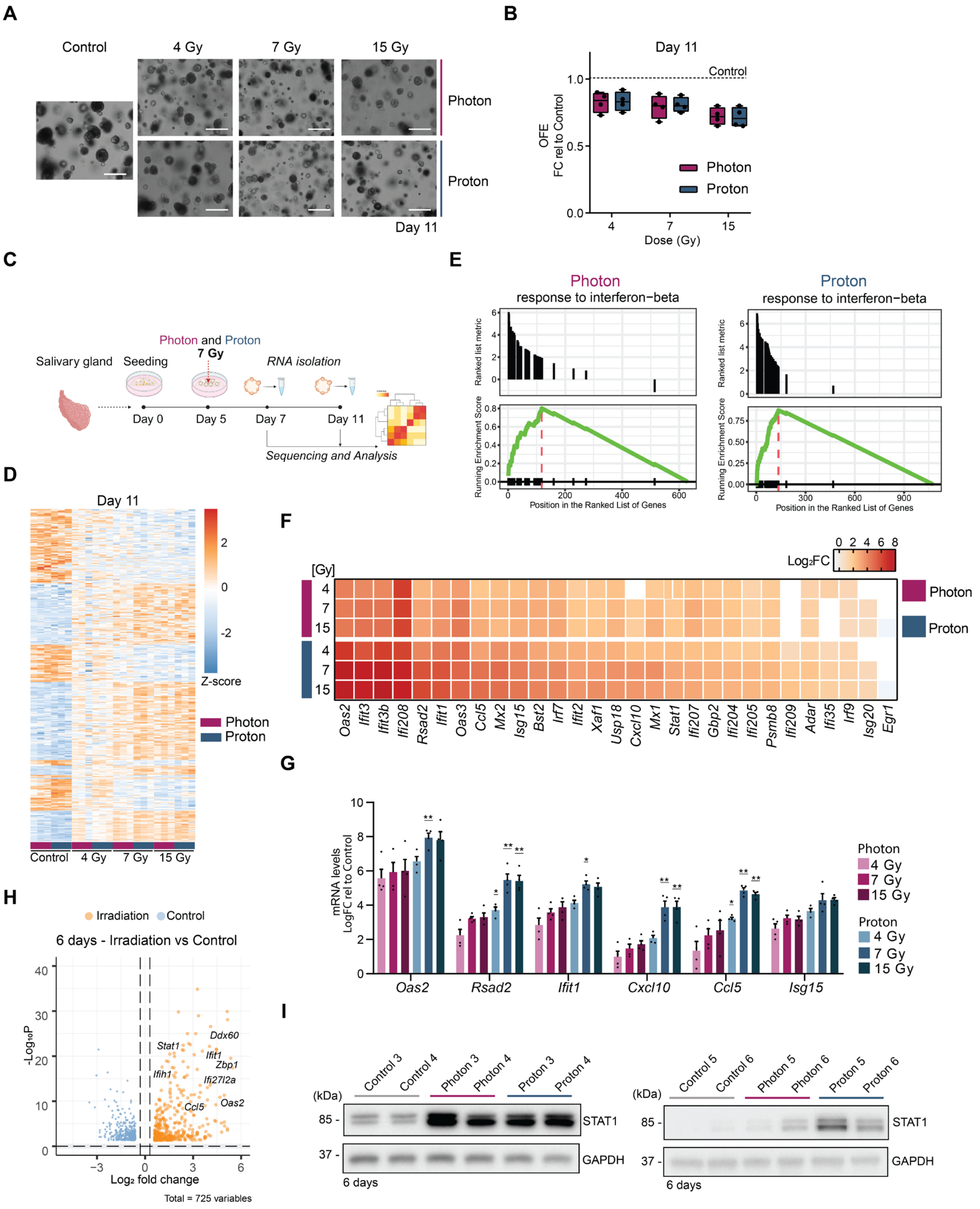


**Fig. S1. Radiation-induced IFN-I response at 6 days post irradiation.**

(**A**) Representative images of organoids in culture at day 11 (6 days post irradiation). Scale bar, 100 µm. (**B**) Number of organoids at 6 days post photon and proton irradiation. Data is shown as FC relative to control (0 Gy). n = 4 animals/condition. (**C**) Schematic representation of the bulk RNA-seq analysis. RNA was isolated from organoids at 2 days (day 7) and 6 days (day 11) post-irradiation. (**D**) Heatmap extrapolated from bulk RNA-seq analysis showing differences in gene expression (Z-score) between irradiated and control organoids at 6 days post irradiation using different radiation doses (4, 7 and 15 Gy). Gene list and Z-score are provided in supplementary material. (**E**) Enrichment plots of the GO term "response to interferon-beta" at 6 days post irradiation. (**F**) Heatmap extrapolated from bulk RNA-seq data showing ISGs expression at different doses (4, 7 and 15 Gy) at 6 days post irradiation. Data is shown as Log2FC relative to control. (**G**) rt-qPCR analysis of ISGs at different doses (4, 7 and 15 Gy) at 6 days post irradiation. Data is shown as FC relative to control (means ± s.e.m; n = 4 animals/condition). One-way ANOVA with Tukey`s multiple comparison test. *p and **p are relative to photon. (**H**) Volcano plot showing significant ($p < 0.05$) differentially expressed genes at 6 days post irradiation. (**I**) Western blot analysis of STAT1 and GAPDH of organoids at 6 days post irradiation. In all figures *$p < 0.05$, **$p < 0.01$.

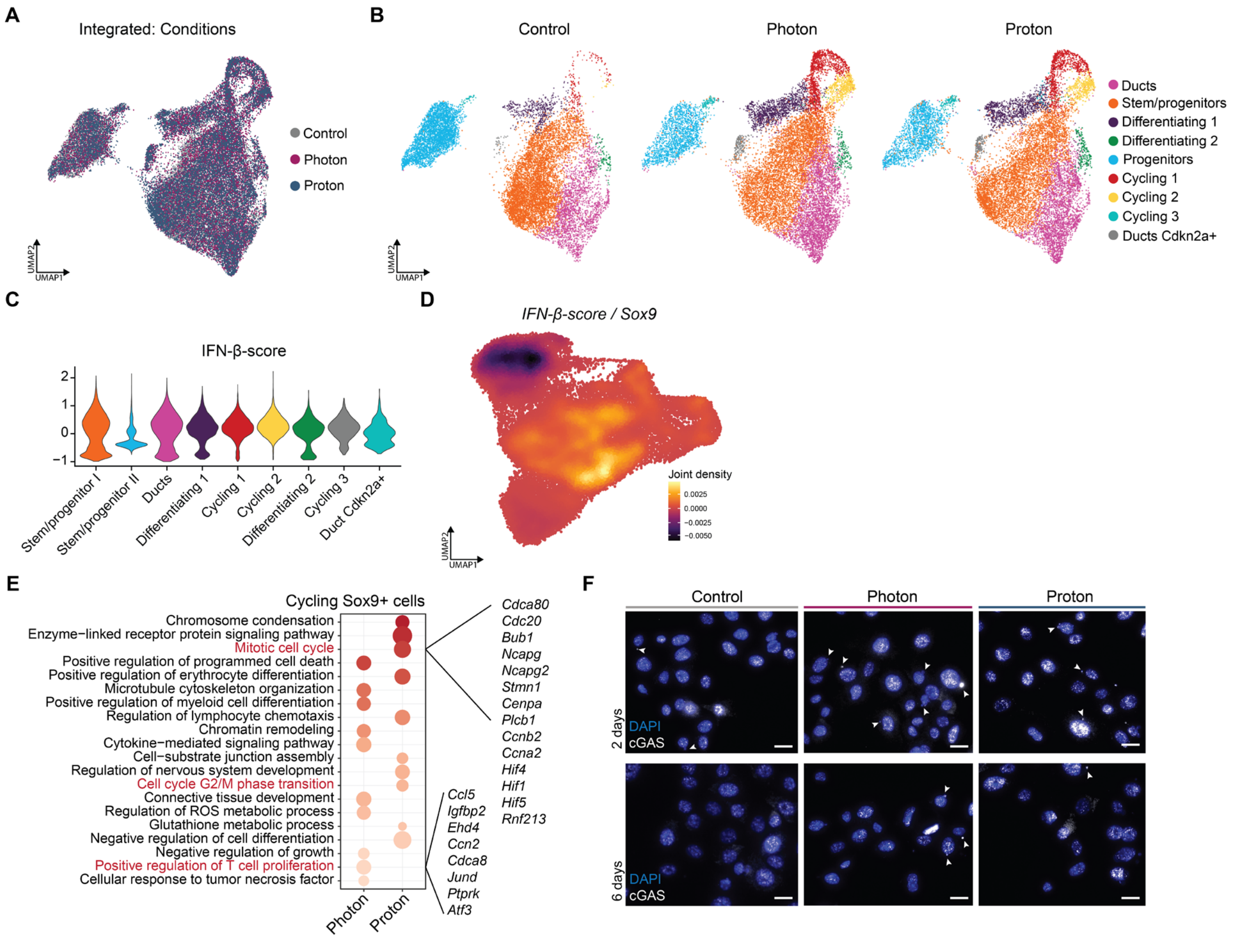


#### Fig. S2. scRNA-seq analysis.

(**A**) UMAP of the integrated dataset of organoids at 6 days post irradiation (day 11) showing the different conditions. (**B**) UMAPs of the integrated dataset of organoids at 6 days post irradiation (day 11) showing the main cell populations in each condition (control, photon and proton). (**C**) Violin plot showing IFN-I-score in each population. (**D**) Density plots showing the joint expression of Sox9 and IFN-I-score in the non-integrated UMAP. (**E**) Top 10 significant (*p*-value < 0.05) upregulated biological processes in proton vs control and photon vs control in the cycling *Sox9*+ cells. Dot color represents *p*-value (-LogPv), dot size represents gene number. Processes related to cell cycle progression are highlighted in red and genes associated to these processes are shown on the right. (**F**) Representative images of immunofluorescence staining of salivary gland cells at 2 days (top panel) and 6 days (bottom panel) post irradiation. Arrows show cGAS+ micronuclei. Scale bar, 5 µm.

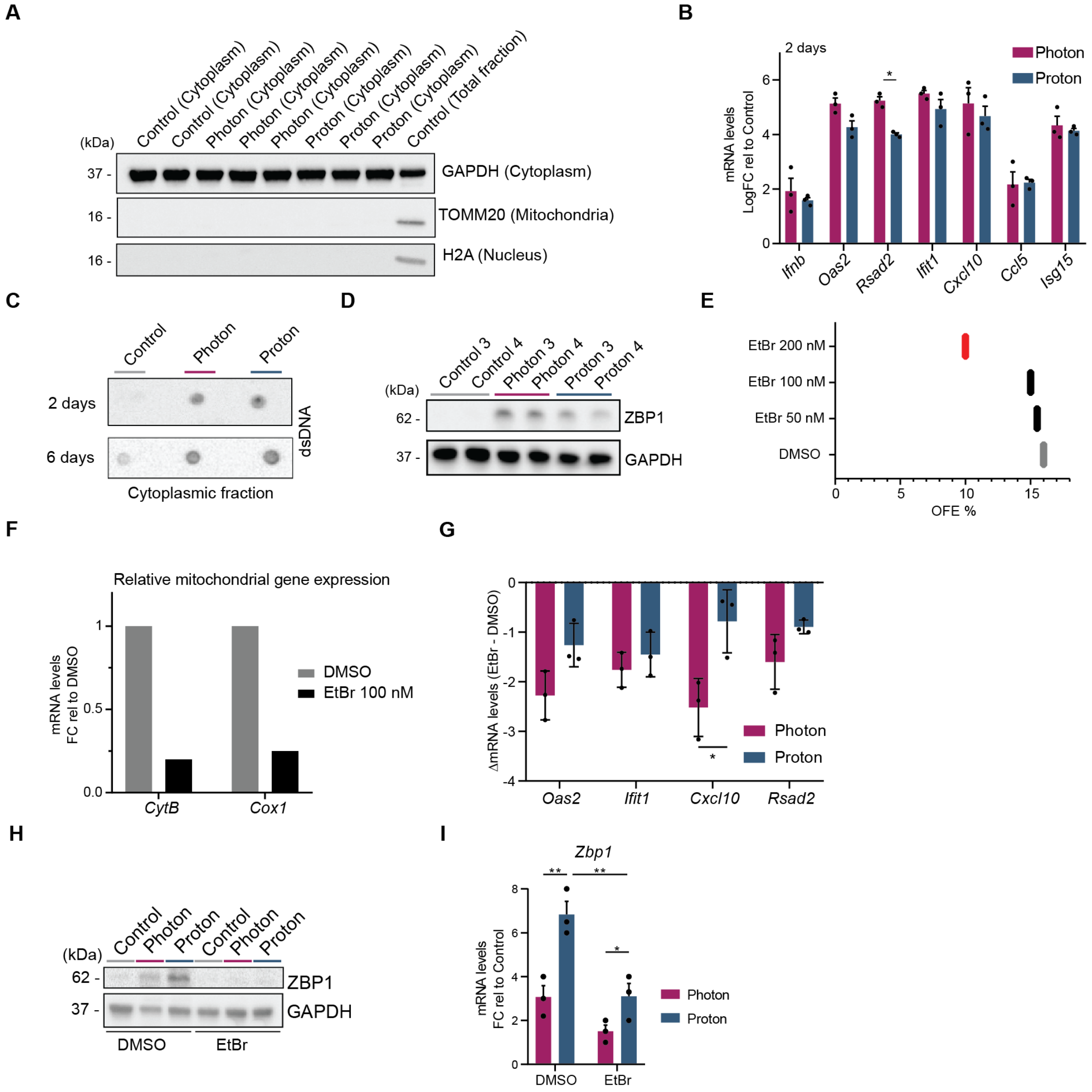

### Fig. S3. Modulation of dsDNA sensing pathway.

(**A**) Validation of the cytoplasmic fractions. Western blot analysis of GAPDH (cytoplasm), TOMM20 (mitochondria) and H2A (nucleus). (**B**) rt-qPCR analysis of ISGs at 2 days post irradiation. Data is shown as FC relative to control (means ± s.e.m; n = 3 animals/condition). Two-sided unpaired *t*-test. (**C**) Dot blot analysis of dsDNA of cytoplasmic fractions extracted from organoids at 2 days and 6 days post irradiation. (**D**) Western blot analysis of ZBP1 and GAPDH at 6 days post irradiation. (**E**) Organoid quantification shown as OFE% showing cytotoxicity after treatment with EtBr 200 nM (red) but not after 50 and 100 nM (black). (**F**) rt-qPCR analysis of mitochondrial DNA genes normalized to nuclear DNA genes showing a decrease of mtDNA content after treatment with EtBr 100 nM at day 11. (**G**) Delta (Δ) values of rt-qPCR shown in Figure 2J (EtBr – DMSO) (means ± s.e.m; n = 3 animals/condition). Two-sided unpaired *t*-test. (**H**) Western blot analysis of ZBP1 and GAPDH in DMSO and EtBr-treated organoids at 6 days post irradiation. (**I**) rt-qPCR analysis of ZP1 in DMSO and EtBr-treated organoids at 6 days post irradiation. (means ± s.e.m; n = 3 animals/condition). One-way ANOVA with Tukey`s multiple comparison test. In all figures *p < 0.05, **p < 0.01.

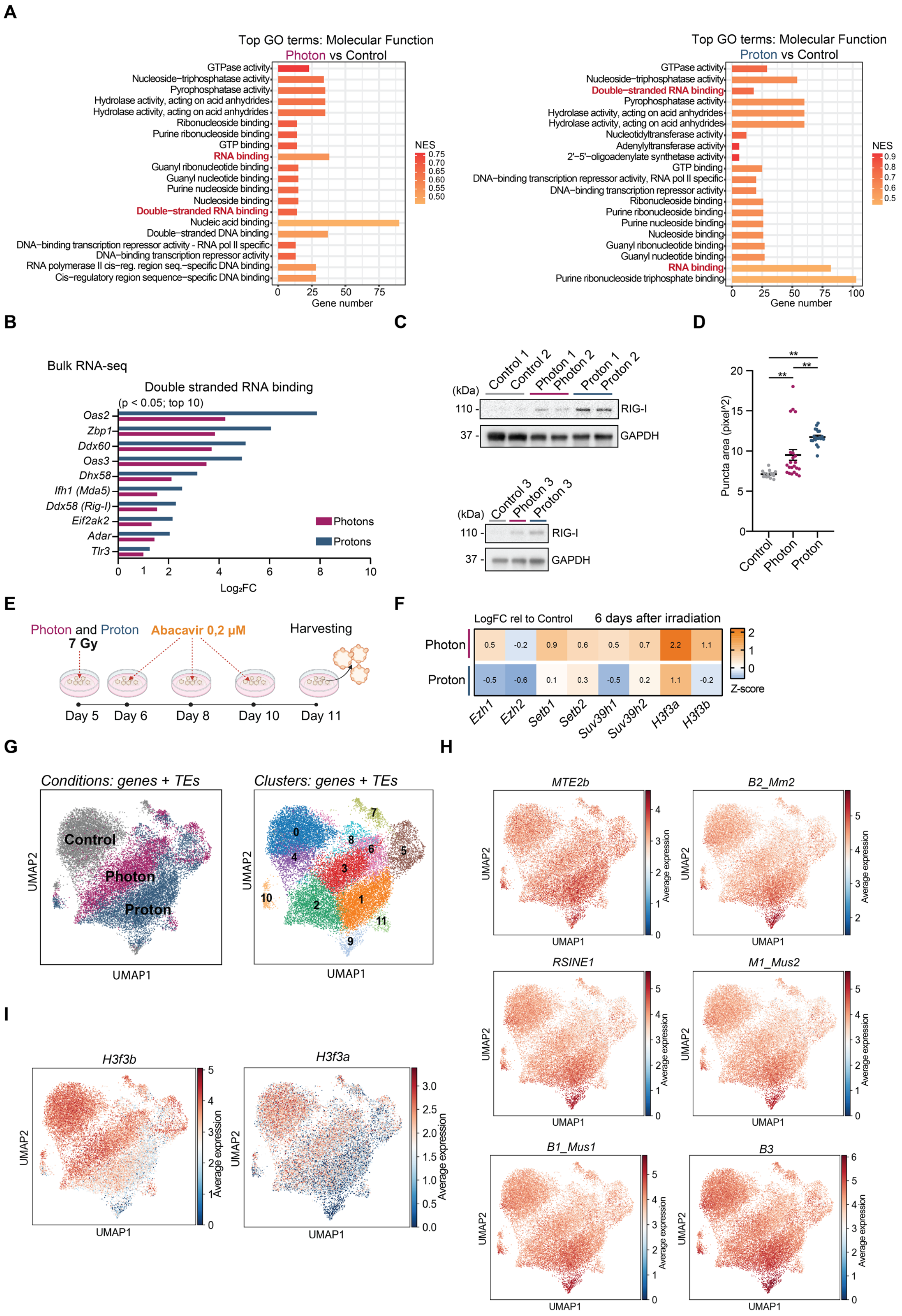
A
Top GO terms: Molecular Function
Photon vs Control
GTPase activity
Nucleoside−triphosphatase activity
Pyrophosphatase activity
Hydrolase activity, acting on acid anhydrides
Hydrolase activity, acting on acid anhydrides
Ribonucleoside binding
Purine ribonucleoside binding
GTP binding
RNA binding
Guanyl ribonucleotide binding
Guanyl nucleotide binding
Purine nucleoside binding
Nucleoside binding
Double−stranded RNA binding
Nucleic acid binding
Double−stranded DNA binding
DNA−binding transcription repressor activity - RNA pol II specific
DNA−binding transcription repressor activity
RNA polymerase II cis−reg. region seq.−specific DNA binding
Cis−regulatory region sequence−specific DNA binding
NES
0.75
0.70
0.65
0.60
0.55
0.50
0
25
50
75
Gene number
Top GO terms: Molecular Function
Proton vs Control
GTPase activity
Nucleoside−triphosphatase activity
Double−stranded RNA binding
Pyrophosphatase activity
Hydrolase activity, acting on acid anhydrides
Hydrolase activity, acting on acid anhydrides
Nucleotidyltransferase activity
Adenylyltransferase activity
2'−5'−oligoadenylate synthetase activity
GTP binding
DNA−binding transcription repressor activity, RNA pol II specific
DNA−binding transcription repressor activity
Ribonucleoside binding
Purine ribonucleoside binding
Purine nucleoside binding
Nucleoside binding
Guanyl ribonucleotide binding
Guanyl nucleotide binding
RNA binding
Purine ribonucleoside triphosphate binding
NES
0.9
0.8
0.7
0.6
0.5
0
25
50
75
100
Gene number
B
Bulk RNA-seq
Double stranded RNA binding
(p < 0.05; top 10)
Oas2
Zbp1
Ddx60
Oas3
Dhx58
Ifh1 (Mda5)
Ddx58 (Rig-I)
Eif2ak2
Adar
Tlr3
Photons
Protons
0
1
2
4
6
8
10
Log₂FC
C
(kDa)
Control 1
Control 2
Photon 1
Photon 2
Proton 1
Proton 2
110 -
RIG-I
37 -
GAPDH
(kDa)
Control 3
Photon 3
Proton 3
110 -
RIG-I
37 -
GAPDH
D
**
**
**
Puncta area (pixel^2)
0
5
10
15
20
Control
Photon
Proton
E
Photon and Proton
7 Gy
Abacavir 0,2 μM
Harvesting
Day 5
Day 6
Day 8
Day 10
Day 11
F
LogFC rel to Control
6 days after irradiation
Photon
0.5
-0.2
0.9
0.6
0.5
0.7
2.2
1.1
Proton
-0.5
-0.6
0.1
0.3
-0.5
0.2
1.1
-0.2
2
1
0
Z-score
Ezh1
Ezh2
Setb1
Setb2
Suv39h1
Suv39h2
H3f3a
H3f3b
G
Conditions: genes + TEs
Control
Photon
Proton
UMAP2
UMAP1
Clusters: genes + TEs
0
1
2
3
4
5
6
7
8
9
10
11
UMAP2
UMAP1
H
MTE2b
B2_Mm2
RSINE1
M1_Mus2
B1_Mus1
B3
UMAP2
UMAP1
Average expression
I
H3f3b
H3f3a
UMAP2
UMAP1
Average expression
3.0
2.5
2.0
1.5
1.0
0.5
0.0

**Fig. S4. Upregulation of TEs and dsRNA-related genes after irradiation.**

(**A**) Bar blot showing the significant top 20 upregulated gene ontology terms related to molecular function. Photon (left) and proton (right) irradiation lead to enrichment of the term double-stranded RNA binding (in red) at 6 days post-irradiation. (**B**) ) Bulk RNA-seq analysis of the significant top 10 genes related to the gene ontology term double-stranded RNA binding. Data extrapolated from bulk RNA-seq analysis shown as $Log_2FC$ relative to control. (**C**) Western blot analysis of RIG-I at 6 days post irradiation (western blot of samples 1 and 2 is the same as in Fig. 3C). (**D**) Immunofluorescence staining quantification (Fig. 3F) showing the puncta mean area (means ± s.e.m; dots represent number of organoids counted/condition from 3 different animals). One-way ANOVA with Tukey`s multiple comparison test. (**E**) Schematic representation of Abacavir treatment. Irradiated organoids were treated every 2 days and harvested on day 11 for analysis. (**F**) Gene expression analysis of heterochromatin regulators at 6 days post irradiation. Data extrapolated from bulk RNA-seq analysis shown as $Log_2FC$ relative to control. (**G**) UMAPs of the non-integrated dataset of organoids at 6 days post irradiation (day 11) re-analyzed considering both genes and TEs showing conditions (top) and main clusters (bottom). (**H**) UMAPs of the re-analyzed non-integrated dataset of organoids at 6 days post irradiation (day 11) showing the expression of different TEs considered for TE-score*: MTE2b, B2_Mm2, RSINE1, M1_Mus2, B1_Mus1* and *B3*. (**I**) UMAPs of the re-analyzed non-integrated dataset of organoids at 6 days post irradiation (day 11) showing the expression of *H3f3b* (top) and *H3f3a* (bottom). In all figures $*p < 0.05$, $**p < 0.01$.

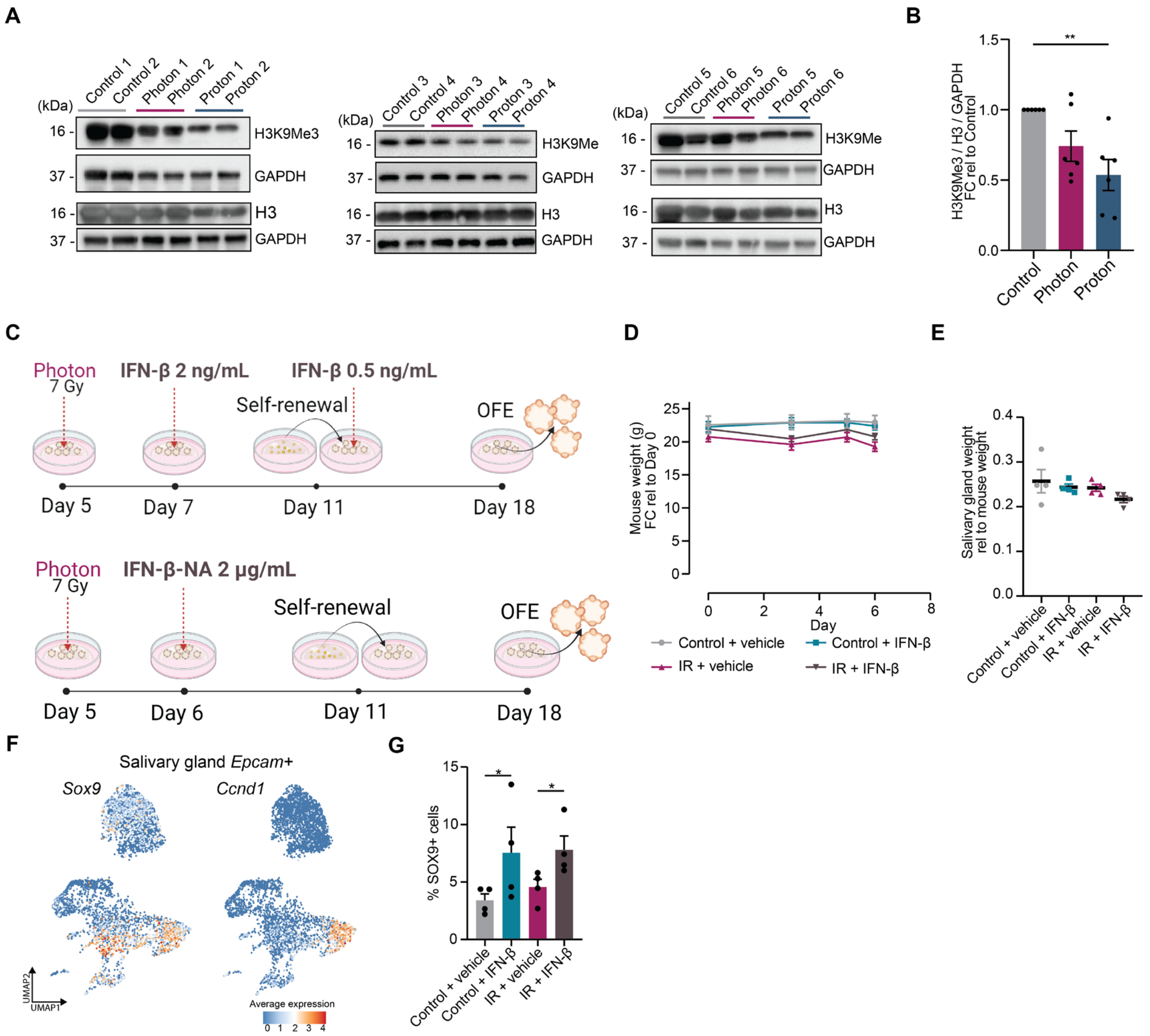


**Fig. S5. Response to IFN-β treatment and irradiation.**

(**A**) Western blot analysis of H3K9Me, H3 and GAPDH of organoids at 6 days post irradiation (western blot of H3K9Me3/GAPDH of mouse 1 and 2 is the same as Fig. 3L). (**B**) Quantification of the western blots in Figure S4B. H3K9Me3 protein levels were normalized for H3 and GAPDH. Data is shown as FC relative to control (means ± s.e.m; n = 6 animals/condition). One-way ANOVA with Tukey`s multiple comparison test. (**C**) Schematic representation of IFN-β treatment (top) and IFN-β-NA treatment (bottom). Organoids were harvested after self-renewal (day 18) for the analysis. (**D**) Analysis of mouse weight over time (means ± s.e.m; n = 4 animals/condition). (**E**) Analysis of salivary gland weight normalized for mouse weight (means ± s.e.m; n = 4 animals/condition). (**F**) UMAP of the subpopulation of *Epcam*+ cells extrapolated from the scRNA-seq dataset of salivary gland tissue [E-MTAB-13374] showing *Sox9* and *Ccnd1* average expression. (**G**) Immunofluorescence staining quantification (Fig. 4H) showing the percentage (%) of SOX9+ cells (means ± s.e.m; n = 4 animals/condition). Two-sided unpaired t-test.

| Reagents and antibodies | Company | Catalog number |
|---|---|---|
| **Organoid culture and self-renewal assay** | | |
| DMEM/F12 | Gibco/Invitrogen | 11320-074 |
| Penicillin- streptomycin antibiotics | Invitrogen | 15140-163 |
| Glutamax | ThermoFisher Scientific | 35050038 |
| EGF | Sigma-Aldrich | E9644 |
| FGF2 | Peprotech | 100-18-B |
| N2 | Gibco | 17502-048 |
| Insulin | Sigma-Aldrich | I6634-100MG |
| Dexamethasone | Sigma-Aldrich | d4902-25mg |
| 0.05% trypsin EDTA | Invitrogen | 25300-096 |
| BME | R&D systems | 3532-010-02P |
| Matrigel | Vwr | 356235 |
| Y27632 | Abcam | ab120129 |
| **Organoid treatments** | | |
| Ethidium Bromide | Sigma | E-4391 |
| C-176 | MedChemExpress | HY-112906 |
| IFN-beta Protein | MedChemExpress | HY-P73130 |
| BRD4770 | MedChemExpress | HY-16705 |
| Mouse IFN-beta Antibody (NA) | R&D systems | MAB8234-100 |
| **Bulk RNA sequencing** | | |
| RNeasy Mini Kit | Qiagen | 74104 |
| High Sensitivity RNA ScreenTapes | Agilent | 5067-5579 |
| QuantSeq 3' mRNA-Seq Library Prep Kit | Lexogen | 015.96 |
| **Western blot** | | |
| STAT1 (1:1000) | Cell Signaling | 14994T |
| GAPDH (1:10000) | Fitzgerald | 10R-G109a |
| cGAS (1:1000) | Cell Signaling | 31659S |
| phospho-STAT1 (Tyr701) (1:1000) | Cell Signaling | 9167S |
| ZBP1 (1:1000) | AdipoGen Life Sciences | AG-20B-0010-C100 |
| dsRNA (1:500) | Scicons | 10020200 |
| RIG-I (1:1000) | Santa Cruz Biotechnology | sc-376845 |
| Tri-Methyl-Histone H3 (Lys9) (1:1000) | Cell Signaling | 13969T |
| H3 (1:5000) | Cell Signaling | 9715S |
| TOMM20 (1:1000) | Abcam | ab78547 |
| H2A (1:1000) | Abcam | ab18255 |
| ECL anti-rabbit IgG HRP | GE-Healthcare | NA934 |
| ECL anti-mouse IgG HRP | GE-Healthcare | NXA931V |
| **Quantitative real-time qPCR** | | |
| dNTP mix | Invitrogen | 10297-018 |

| | | |
|---|---|---|
| Random primers | Invitrogen | SO142 |
| First-strand Buffer | Invitrogen | 28025013 |
| DTT | Invitrogen | 328025013 |
| RNase OUTTM | Invitrogen | 10777019 |
| M-MLV RT | Invitrogen | 28025013 |
| iQ SYBR Green Supermix | Bio-Rad | 170-8885 |
| **Immunofluorescence staining** | | |
| cGAS (1:500) | Cell Signaling | 31659S |
| dsRNA (1:500) | Scicons | 10020200 |
| SOX9 (1:200) | Cell Signaling | 82630T |
| Tri-Methyl-Histone H3 (Lys9) (1:500) | Cell Signaling | 13969T |
| DAPI | Sigma Aldrich | D9542 |
| Alexa Flour 594 goat anti-rabbit (1:1000) | ThermoFisher Scientific | A-11012 |
| Opal 4-Color anti-Rabbit Manual IHC Kit | Akoya | NEL810001KT |
| **Subcellular fractionation** | | |
| Sucrose | Sigma Aldrich | 57-50-1 |
| HEPES | Sigma Aldrich | H3784-100G |
| KCl | Merck | 7447-40-7 |
| $MgCl_2$ | Sigma Aldrich | 7786-30-3 |
| EDTA | Invitrogen | 6381-92-6 |
| EGTA | Sigma Aldrich | 67-42-5 |
| DTT | Sigma Aldrich | D0632-5G |
| **Dot blot** | | |
| dsDNA (1:1000) | Santa Cruz Biotechnology | HYB331-01 |
| **Flow cytometry** | | |
| TMRE | MedChemExpress | HY-D0985A |
| Mitotracker Green | ThermoFisher Scientific | M7514 |
| **Single cell RNA-sequencing** | | |
| 10X Chromium Next GEM Single Cell 3' Kit v3.1 | 10X Genomics | 1000128 |
| Single Index Kit T Set A | 10X Genomics | 1000213 |
| 10X Chromium Next GEM Chip G Single Cell Kit | 10X Genomics | 1000127 |

**Table S1. Details of all reagents and antibodies used in this study.**

| Gene | Forward Primer 5`- 3` | Reverse Primer 5` - 3` |
|---|---|---|
| *Ywhaz* | TTACTTGGCCGAGGTTGCT | TGCTGTGACTGGTCCACAAT |
| *Oas2* | TTGAAGAGGAATACATGCGGAA | GGGTCTGCATTACTGGCACTT |
| *Rsad2* | GGTGCCTGAATCTAACCAGAAG | CCACGCCAACATCCAGAATA |
| *Ifit1* | CTGAGATGTCACTTCACATGGAA | GTGCATCCCCAATGGGTTCT |
| *Cxcl10* | GCCGTCATTTTCTGCCTCA | CGTCCTTGCGAGAGGGATC |
| *Ccl5* | TGCCCACGTCAAGGAGTATTTC | AACCCACTTCTTCTCTGGGTTG |
| *Isg15* | GGTGTCCGTGACTAACTCCAT | TGGAAAGGGTAAGACCGTCCT |
| *Ddx58 (Rig-I)* | AAGAGCCAGAGTGTCAGAATCT | AGCTCCAGTTGGTAATTTCTTGG |
| *Ifih1 (Mda5)* | GTGATGACGAGGCCAGCAGTTG | ATTCATCCGTTTCGTCCAGTTTC A |
| *Zbp1* | AAGAGTCCCCTGCGATTATTTG | TCTGGATGGCGTTTGAATTGG |
| *IAPEz-int* | AGGATGAAGACCCCCACCTT | AGAAGCAGAAGGAGCAGCAG |
| *IAPEY3-A* | AGGAGGATTGGAGTTTTGGCA | ACTTCTTCACGTAGAAATGCTCT |
| *RLTR4* | AGGTGACCTAACCCTCCTCC | GCGACCTACAAAGAGGACTCA |
| *MERVL-int* | ATACCACAAAGAACTCAAGGGCA | TGGAATTGATGCGAAGCTCCC |
| *L1Md-A* | AGGAGCTTGCTGCTCACTTT | ATTCGTGCCACTCCTGTCTC |
| *L1Md-G* | TAAGCAGCTGTCCAGCCAAA | GGCACAGCCCTTGAGTAGAG |
| *mt16S* | CACTGCCTGCCCAGTGA | ATACCGCGGCCGTTAAA |
| *mtCOX2* | CTACAAGACGCCACAT | GAGAGGGGAGAGCAAT |
| *mtDloop3* | TCCTCCGTGAAACCAACAA | AGCGAGAAGAGGGGCATT |

**Table S2. Primer sequences.**
